# CODANIN-1 sequesters ASF1 by using a histone H3 mimic helix to regulate the histone supply

**DOI:** 10.1101/2024.07.10.602876

**Authors:** Tae-Kyeong Jeong, R. Ciaran MacKenzie Frater, Jongha Yoon, Anja Groth, Ji-Joon Song

## Abstract

ASF1 is a major histone chaperone that regulates the supply of histone H3–H4 and facilitates nucleosome assembly to maintain chromatin structure during DNA replication and transcription. CODANIN-1 negatively regulates the function of ASF1. However, the molecular mechanism by which CODANIN-1 inhibits the ASF1-mediated histone supply remains elusive. Here, we present the electron microscopy (cryo-EM) structure of a human CODANIN-1_ASF1A complex at 3.75 Å resolution. The structure reveals that CODANIN-1 forms a dimer where each monomer holds two ASF1 molecules, utilizing two B-domains and two histone H3 mimic helices (HMHs). The interaction of CODANIN-1 with ASF1 via the HMH and B domains inhibits the formation of an ASF1/H3–H4 complex and sequesters ASF1 in the cytoplasm. Our study provides a structural and molecular basis for the function of CODANIN-1 as a unique negative regulator that highjacks ASF1 interaction sites with histones and downstream chaperones to inhibit nucleosome assembly.

## Introduction

Eukaryotic DNA is organized into a dynamic multilayered structure called chromatin. The regulation and maintenance of chromatin structure are essential for expressing and safeguarding genomic information. The fundamental repeating unit of chromatin is the nucleosome, which consists of two copies of each of the four core histones (H3, H4, H2A and H2B) and 147 base pairs of DNA^1^. The nucleosome is a dynamic structure and the major target of chromatin regulation. The nucleosome position and structure are remodeled in an ATP-dependent manner, and histones are covalently modified with various moieties, including methyl, acetyl, phosphoryl, and ubiquitin moieties^2–4^. The combination of nucleosome remodeling and histone modifications governs genome regulation. Nucleosomes are dynamically disassembled and assembled to maintain chromatin integrity during cellular processes such as DNA replication and transcription^5–7^. The assembly of nucleosomes occurs in a stepwise manner, assisted and regulated by a group of proteins called histone chaperones. Histone chaperones shield histones from unwarranted interactions and facilitate their delivery to DNA and assembly into nucleosomes. Anti-silencing function 1 (ASF1) is a major histone H3‒H4 chaperone in the cytoplasm and nucleus^5,8^ and regulates the supply of histones to sites of ongoing transcription and replication. In mammals, there are two ASF1 isoforms (ASF1A and ASF1B), which show a high level of sequence conservation except in the unstructured C-terminal region. While ASF1A is ubiquitously expressed, ASF1B is expressed mainly in proliferating cells^9^. In the cytoplasm, newly synthesized histone H3‒H4 dimers are formed by DNAJC9 and nuclear autoantigenic sperm protein (NASP) before they are modified by the HAT1_RbAP46/48 histone acetyltransferase complex at lysines 5 and 12 on histone H4^10–14^. ASF1, in turn, engages histone H3–H4 in complex with NASP and mediates the nuclear import of histone H3–H4 by importins^10,11,15,16^. In the nucleus, ASF1 delivers histone H3–H4 dimers to CAF-1^17,18^ and the HIRA^19–21^ complexes for nucleosome assembly during replication-dependent and replication-independent processes, respectively. Thus, ASF1 directly regulates the flow of newly synthesized histone H3–H4 to chromatin across the cell cycle, with its major task involving the histone supply for DNA replication in S phase ^8,22–24^. ASF1 directly interacts with the αH3 helix of histone H3, part of the histone H3–H4 tetramerization interface, and prevents histone H3–H4 from forming a tetramer prior to nucleosome assembly^25,26^. To hand over histones to downstream chaperones, ASF1 interacts with the B-domains in CAF-1 and HIRA using an interface on the opposite side of its histone-binding surface^27,28^. We previously reported that CODANIN-1 interacts with ASF1 via a B domain and negatively regulates the function of ASF1^29^. CODANIN-1 overexpression sequesters ASF1 in the cytoplasm and induces cell cycle arrest in S phase, indicating that CODANIN-1 inhibits the ASF1-mediated histone supply. CODANIN-1, which is encoded by *CDAN1,* forms a complex with CDAN1-interacting nuclease 1 (CDIN1)^30–32^. Like ASF1A, CODANIN-1 and CDIN1 are ubiquitously expressed, and mutations in CDAN1 or CDIN1 can cause the rare, autosomal recessive disease congenital dyserythropoietic anemia type I (CDA-I)^33,34^. CDA-I is characterized by defective terminal erythropoiesis and nuclear abnormalities such as a spongy appearance of heterochromatin in erythroblasts ^35–37^. While the exact functions of CODANIN-1 and CDIN1 in erythropoiesis are unknown, their interaction with ASF1 suggests that the regulation of histone dynamics might be critical for chromatin compaction in the late stages of erythropoiesis. In this study, we provide a structural and molecular basis for the mechanism by which CODANIN-1 sequesters and inactivates ASF1.

## Results

### Cryo-EM structure of the CODANIN-1 complex

To understand how CODANIN-1 regulates ASF1 function, we determined the cryo-EM structure of the CODANIN-1 complex, which is composed of full-length CODANIN-1 (1-1,227 a.a.), ASF1A (1-172 a.a.) and full-length CDIN1 (a.k.a. C15ORF141) (1-281 a.a.) (Fig. 1a). The CODANIN-1 complex was estimated to be more than 400 kDa by size-exclusion chromatography (Fig. 1b), suggesting that CODANIN-1 forms a multimer. For the cryo-EM study, we further purified the CODANIN-1 complex with GraFix and collected 4,282 micrographs using a Titan Krios with a Gatan K3 detector. We then processed the data, generating a 3.75 Å resolution structure of the CODANIN-1 complex (Figs. 1c, 1d, Supplementary Fig. 1 and Table 1). The high quality of the cryo-EM map with the help of AlphaFold 2 enabled us to generate an atomic model of the CODANIN-1 complex (Figs. 1e, 1f, and Supplementary Fig. 2). The C-terminal region of CODANIN-1 and the entire CDIN1 region were not visible in the cryo-EM map. This finding likely reflects that both are highly flexible, as CDIN1 is known to bind the flexible C-terminal domain of CODANIN-1^30–32^.

**Fig. 1.**
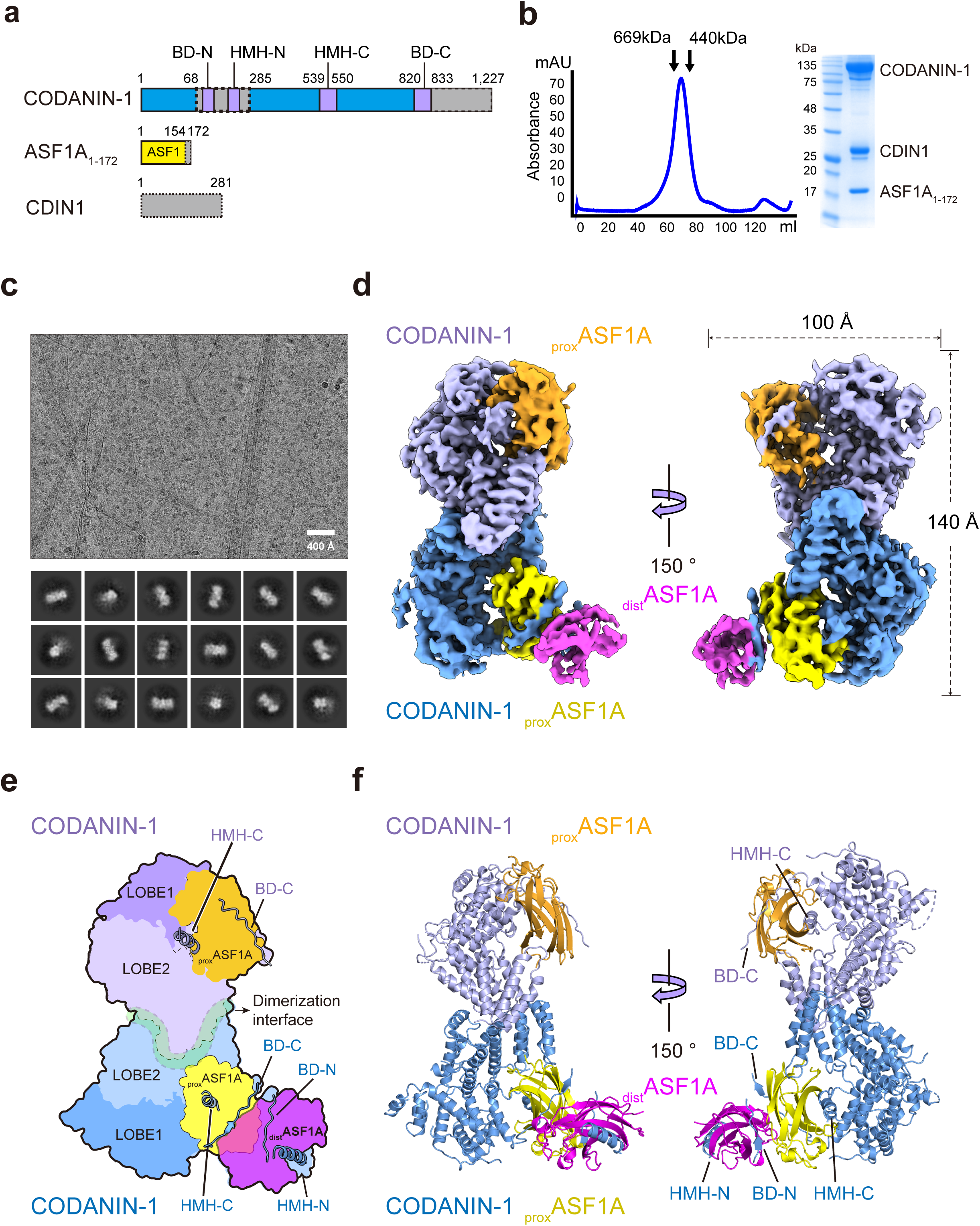
Cryo-EM structure of the CODANIN-1 complex. **a,** Schematic diagram of the CODANIN-1 complex composed of CODANIN-1 full-length (1-1,227), C-terminal-truncated ASF1A_1-172_ (1-172 a.a.) and full-length CDIN1 (1-281 a.a.). The dotted boxes indicate the disordered region in the cryo-EM map. **b**, Gel filtration profile and SDS‒PAGE analysis of the CODANIN-1 complex. The gel filtration profile is representative of n=4 biological replicates. **c**, Representative cryo-EM micrograph of the CODANIN-1 complex on a graphene oxide-covered grid and 2D class averages. The micrograph is representative of total 4,075 selected micrographs. **d**, Cryo-EM map of the CODANIN-1 complex. Two CODANIN-1 molecules in a dimer are colored pale blue and pale purple, the bound proximal ASF1A is yellow and orange, and the distal ASF1A is magenta. **e,** Schematic diagram of the structure of the CODANIN-1_ASF1A complex. **f**, The atomic structure of the CODANIN-1_ASF1A complex is shown in a ribbon diagram.

**Table 1.**
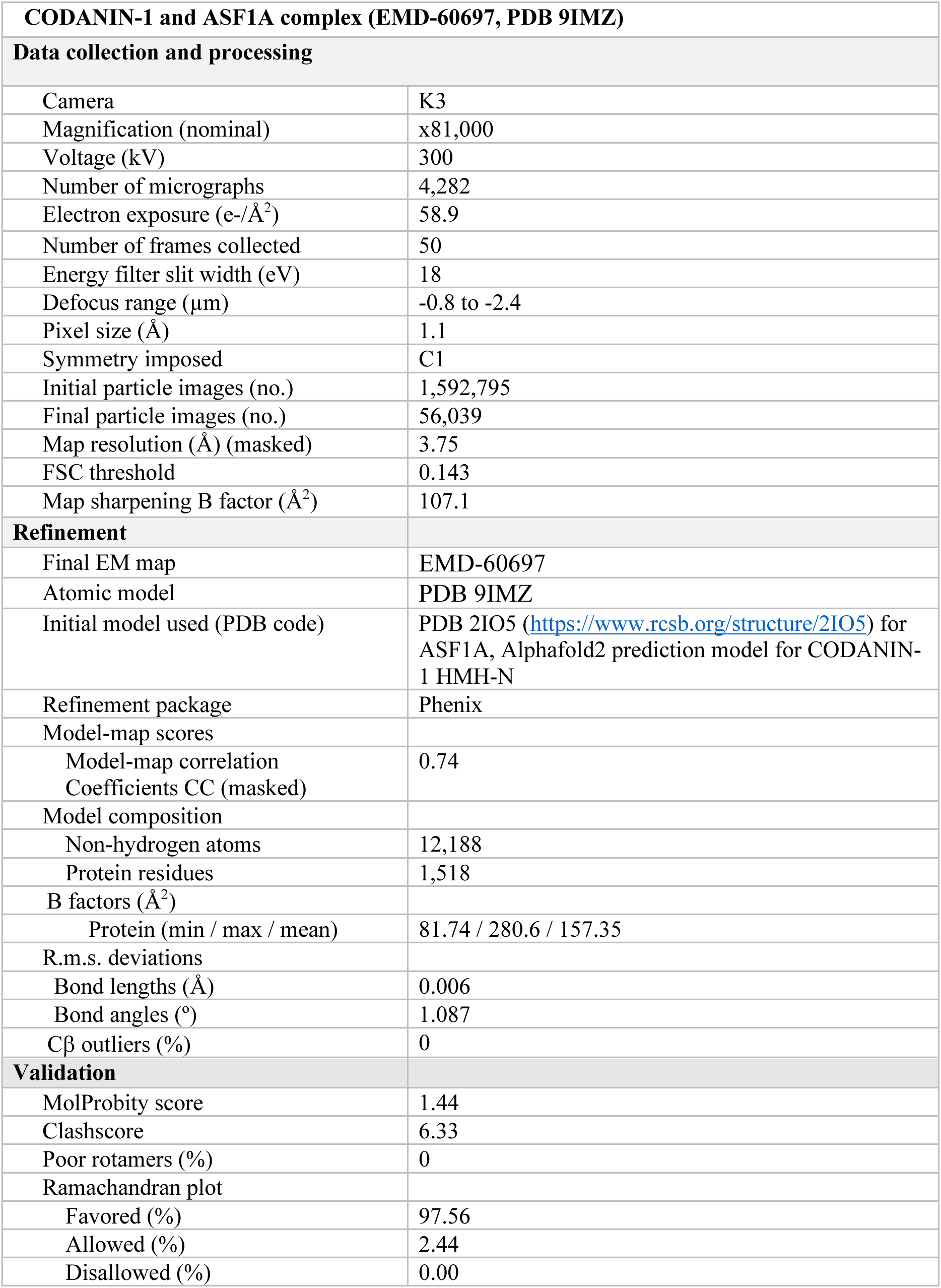
Cryo-EM data collection and processing statistics.

The cryo-EM structure shows that CODANIN-1 dimerizes with each monomer, binding two ASF1A molecules in a proximal (_prox_ASF1A) and distal (_dist_ASF1A) manner (Figs. 1e, 1f and Supplementary Fig. 3). While one of the CODANIN-1 monomers clearly has EM densities for two ASF1 molecules, the other CODANIN-1 monomer clearly has EM density for _prox_ASF1A but a weak density for _dist_ASF1A. However, lowering the threshold of the map enables visualization of the position of _dist_ASF1A (Supplementary Fig. 3), indicating that a monomer can accommodate two ASF1A molecules. At this time, we do not have a clear explanation for why one of the _prox_ASF1A molecues exhibits a poor density. It is possible that the relative orientation between two _dist_ASF1A molecules is very dynamic, leading to one of the _dist_ASF1A molecules being disordered. For clarity, we modeled _dist_ASF1A for only one of the CODANIN-1 dimers (Fig. 1f).

This configuration of the CODANIN-1 dimer associated with multiple ASF1 molecules is consistent with the size of the complex estimated by size-exclusion chromatography (Fig. 1b). CODANIN-1 consists mostly of α-helices (Fig. 1f and Supplementary Figs. 4, 5), which are linked by disordered loops. Specifically, most of the large loops (69-284 a.a.) linking αH4 and αH5 are disordered in the cryo-EM map. In addition, the region beyond 833 a.a. is not visible in the cryo-EM structure, although the C-terminal region is predicted to be composed of α-helices. CODANIN-1 is composed of two lobes: α-helices from αH1 to αH12 along with αH28 form one lobe (LOBE1), and helices from αH13 to αH27 (except αH16) form the other lobe (LOBE2) (Fig. 1e and Supplementary Fig. 5). Interestingly, αH16 is located between the two lobes and interacts with _prox_ASF1A. LOBE2 from the two CODANIN-1 molecules extensively interact with each other to form a dimer with a dimerization area of 1,844 Å^2^ for each CODANIN-1. This large dimerization interface implies that CODANIN-1 forms an obligatory dimer. Overall, the cryo-EM structure of the CODANIN-1_ASF1A complex delineates the structural features of CODANIN-1, which tethers multiple ASF1A molecules.

### CODANIN-1 binds multiple ASF1A molecules via B-domains and histone mimic helices (HMHs)

Our cryo-EM structure delineates the ASF1 binding mode of CODANIN-1. Two ASF1A molecules (_prox_ASF1A and _dist_ASF1A) are associated with one CODANIN-1 subunit in a back-to-back manner, with two β strands sandwiched between the two ASF1A molecules and two separate helices (αH5 for _dist_ASF1A and αH16 for _prox_ASF1A) interacting with their histone H3 binding interface (Figs. 2a and 2f). The two β-strands are B-domains (BDs) that interact with the B-domain binding site of each ASF1A. The β-strand interacting with _dist_ASF1A corresponds to the previously identified B-domain (BD-N:193-202 a.a.) (Fig. 2b)^29^, and the β-strand interacting with _prox_ASF1A is a new second B-domain (BD-C:820-833 a.a.) in CODANIN-1 (Fig. 2c).

**Fig. 2.**
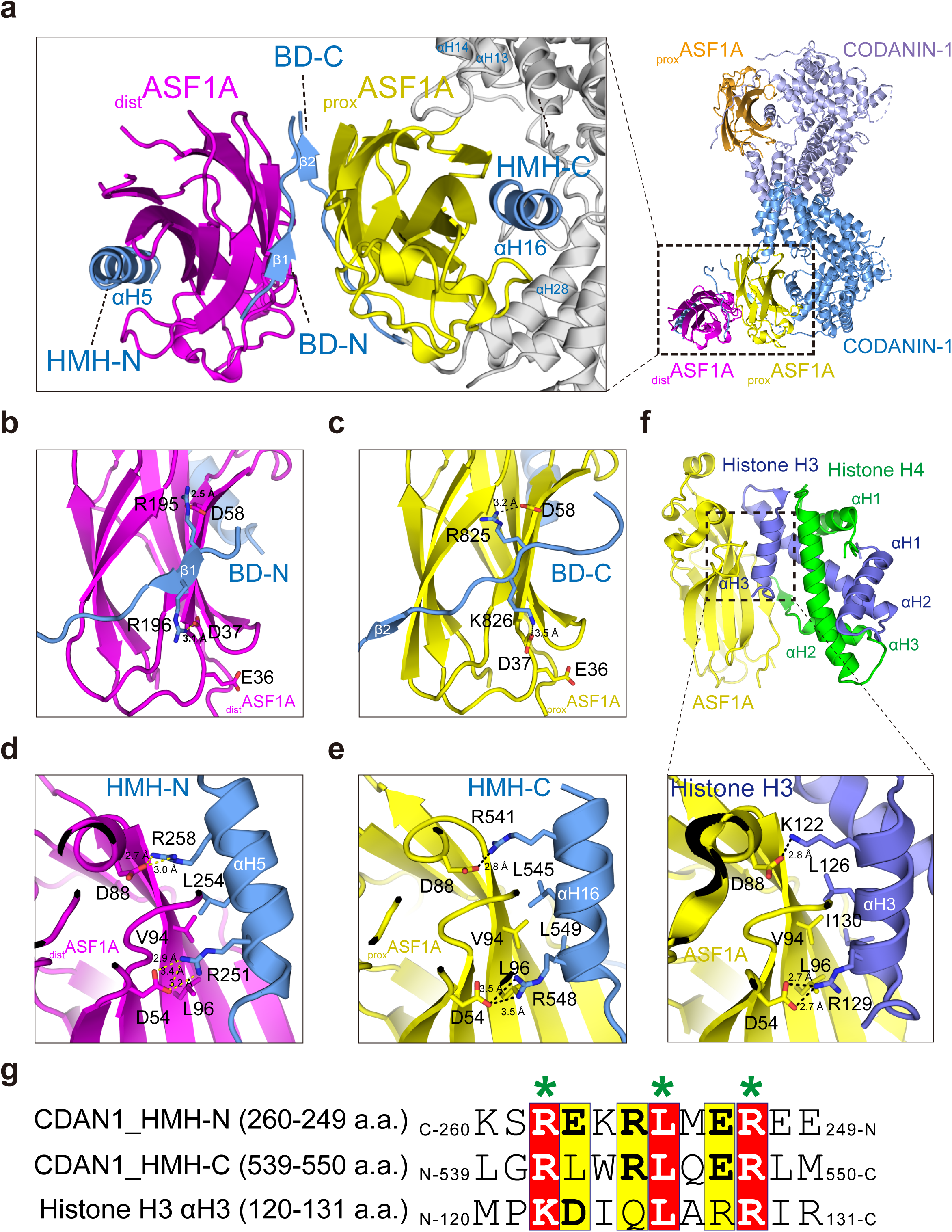
CODANIN-1 has two ASF1A binding modules, each of which is composed of a B-domain (BD) and a histone mimic helix (HMH) **a,** CODANIN-1 forms a dimer, and each molecule binds two ASF1A molecules: proximal ASF1 (_prox_ASF1A) in yellow or orange and distal ASF1A (_dist_ASF1A) in magenta. _prox_ASF1A is encapsulated by BD-C, the αH28 helix, a loop linking the αH13 and αH14 helices, and the αH16 helix (HMH-C). _dist_ASF1A is sandwiched by BD-N and the αH5 helix (HMH-N). **b,c,** Detailed interactions between _dist_ASF1A and BD-N (193-202 a.a.) (b) and between _prox_ASF1A and BD-C (820-833 a.a.) (c). D37 and D58 in _dist_ASF1A (b) and _prox_ASF1A (c) form salt bridges with R196 and R195 in BD-N and K826 and R825 in BD-C, respectively. **d, e, f** Interaction modes of HMH-N with _dist_ASF1A (d), HMH-C with _prox_ASF1A (e) and with histone H3 and the ASF1A complex (PDB 2IO5) (f). HMH-N and HMH-C occupy the histone H3 αH3-binding pockets of ASF1A. V94 in _dist_ASF1A and _prox_ASF1A form hydrophobic interactions with HMH-N L254 and with HMH-C L545 and L549, respectively. D88 and D54 in _dist_ASF1A and _prox_ASF1A form salt bridges with HMH-N R258 and R251 and with HMH-C R541 and R548, respectively. **g**. Sequence comparison between the two CODANIN-1 HMHs (CDAN1_HMH-N and CDAN1_HMH-C) and histone H3 αH3. The residues that interact with ASF1A are marked with asterisks. HMH-N is aligned with HMH-C and histone H3 αH3 in the C-to-N direction.

More specifically, _prox_ASF1A is wrapped by several regions of CODANIN-1, including BD-C, the αH28 helix, a loop linking the αH13 and αH14 helices, and the αH16 helix. Thus, CODANIN-1 essentially encapsulates _prox_ASF1A (Fig. 2a). In contrast, _dist_ASF1A is stacked on _prox_ASF1A via weak electrostatic interactions between the two ASF1A molecules and held by BD-N (193-202 a.a.) and the αH5 helix. Although the two B-domains are sandwiched between _prox_ASF1A and _dist_ASF1A, there is no direct contact between BD-N and BD-C. BD-N (193-202 a.a.) is in the middle of a large loop (69-284 a.a.), which is completely disordered except for the B-domain and the αH5 helix tethered to _dist_ASF1A (Fig. 2b and Supplementary Fig. 5). Therefore, this loop, including BD-N, is likely disordered when CODANIN-1 is not bound to a _dist_ASF1A molecule. The other B-domain, BD-C, is found after the αH28 helix and is separated by a short linker (Supplementary Figs. 4 and 5). The B-domains of CODANIN-1 interact with ASF1A in a similar manner as the B domains in the downstream chaperones CAF-1 and HIRA (Supplementary Fig. 6a). R195 and R196 of BD-N make salt bridges with D58 and D37 of _dist_ASF1A, respectively, whereas R825 and K826 of BD-C interact with D58 and D37 of _prox_ASF1A, respectively (Figs. 2b and 2c). Similar salt bridges are also found in HIRA^27^ and CAF-1^28^ B-domain-ASF1 interactions (Supplementary Fig. 6a).

Interestingly, the CODANIN-1 αH5 (249-260 a.a.) and αH16 helices (539-550 a.a.) interact with the histone-binding pockets of _dist_ASF1A and _prox_ASF1A, respectively. This implies that ASF1A binding to CODANIN-1 is mutually exclusive with histone H3–H4 binding (Figs. 2d-f). Moreover, the amino acid sequences of the αH5 and αH16 helices are very similar to the αH3 helix of histone H3 (Fig. 2g), which is the major ASF1 interaction site in H3–H4 dimers ^25,26^. Notably, the αH5 helix sequence aligns with the αH3 helix of histone H3 in the C-to-N direction (Fig. 2g). Therefore, we designated the αH5 helix the ‘histone mimic helix (HMH)-N’ and the αH16 helix HMH-C. The interactions between the HMHs and the two ASF1A molecules are very similar to those between histone H3 and ASF1A (Figs. 2d-f and Supplementary Fig. 6b). Specifically, L254 of HMH-N hydrophobically interacts with ASF1A V94 and L96, and R251 and R258 of HMH-N form salt bridges with D54 and D88 of ASF1A, respectively (Fig. 2d). For HMH-C, L545 and L549 of HMH-C form hydrophobic interactions with V94 and L96 of ASF1A, and R541 and R548 of HMH-C form salt bridges with D88 and D54 of ASF1A (Fig. 2e). As ASF1A V94, D88 and D54 are critical for the interaction with histone H3 (Fig. 2f)^25,26^, CODANIN-1 seems to harness the binding mode of ASF1A and histone H3 to sequester ASF1A with its HMHs. These data explain why a mutation in the ASF1A histone binding site (V94R) disrupted the association with CODANIN-1 in our recent proteomics study of the histone chaperone network^38^.

### The HMH and B-domain of CODANIN-1 participate in ASF1A binding

The cryo-EM structure shows that CODANIN-1 has two ASF1-binding modules and that each module is composed of both a B-domain and an HMH: the distal ASF1A-binding module (BD-N and HMH-N) and the proximal ASF1A-binding module (BD-C and HMH-C). We first asked whether each HMH or B domain is sufficient for ASF1A binding. We generated GST fusions of the BD-N (193-202 a.a.), HMH-N (249-260 a.a.), HMH-C (539-550 a.a.) and BD-C (820-833 a.a.) domains and performed GST pulldown assays for ASF1A binding (Fig. 3a). Among the four domains, BD-N showed the strongest binding, HMH-C marginally bound to ASF1A, and HMH-N and BD-C showed no binding. These data suggest that BD-N and HMH-C may be the major binding domains for _dist_ASF1A and _prox_ASF1A, respectively.

**Fig. 3.**
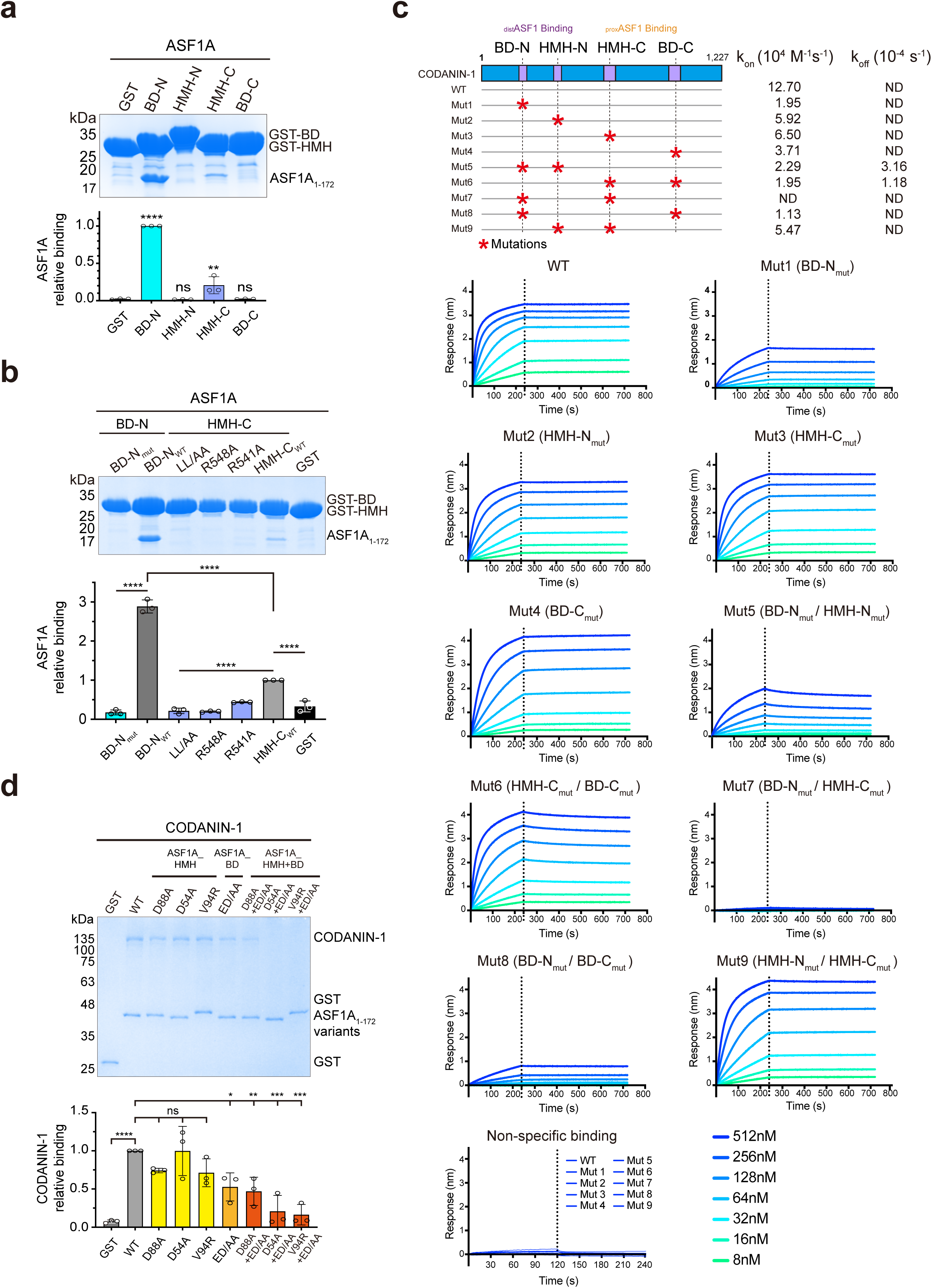
HMH and the B-domain of CODANIN-1 synergistically contribute to ASF1A binding. **a,** GST pulldowns of ASF1A with the GST-BD-N (183-217 a.a.), GST-HMH-N (255-285 a.a.), GST-HMH-C (520-557 a.a.) and GST-BD-N (816-843 a.a.) domains. For the bar graph, each dot represents a biological replicate (n=3), error bars represent the mean±SD. P-values were calculated with one-way analysis of variance (ANOVA) with Dunnett’s multiple comparisons test against GST (BD-N: p<0.0001, HMH-N: p=0.9998, HMH-C: p=0.0043, BD-C: p>0.9999). **b,** GST pulldowns of ASF1A with GST-HMH-C mutants (R541A, R548A or LL/AA) and GST-BD-N mutants (B_mut_-R195A/R196A). For the bar graph, each dot represents a biological replicate (n=3), error bars represent the mean±SD. P-values were calculated with one-way analysis of variance (ANOVA) with Dunnett’s multiple comparisons test (BD-N_mut_: p<0.0001 against BD-N_wt_, LL/AA, R548A and R541A: p<0.0001 against HMH-C_wt_, BD-N_wt_ and HMH-C_wt_: p<0.0001 against GST). **c**, Biolayer interferometry (BLI) measurement of ASF1A binding to full-length CODANIN-1 and the indicated mutants. ASF1A was immobilized on the surface of the chips, and CODANIN-1 was injected at 8, 16, 32, 64, 128, 256, and 512 nM concentrations. The dotted line indicates the time points at which the analytes (CODANIN-1) were removed. Schematic diagram of the mutants (top panel): Mut1 (BD-N_mut_), Mut2 (HMH-N_mut_), Mut3 (HMH-C_mut_), Mut4 (BD-C_mut_), Mut5 (BD-N_mut/_HMH-N_mut_), Mut6 (HMH-C_mut_/BD-C_mut_), Mut7 (BD-N_mut_/HMH-C_mut_), Mut8 (BD-N_mut_/BD-C_mut_) and Mut9 (HMH-N_mut_/HMH-C_mut_). ND (< 10^−7^ or no binding). None of the mutants bound to the chip surface without ASF1A immobilization (nonspecific binding). **d**, GST pulldowns of full-length CODANIN-1 using wild-type GST-ASF1A and the indicated mutants. ASF1A mutants on the HMH binding surface (D88A, D54A, V94R), the B-domain binding surface in ASF1A (ED/AA: E36A/D37A), and double mutants of both (D88A+ED/AA, D54A+ED/AA, V94R+ED/AA) were tested. For the bar graph, each dot represents a biological replicate (n=3), error bars represent the mean±SD. P-values were calculated with one-way analysis of variance (ANOVA) with Dunnett’s multiple comparisons test against WT (GST: p<0.0001, D88A: p=0.3572, D54A: p>0.9999, V94R: p=0.2566, ED/AA: p=0.0209, D88A+ED/AA: p=0.0087, D54A+ED/AA: p=0.0002, V94R+ED/AA: p=0.0001). For the all bar graphs, ns: not significant, *: p < 0.05, **: p < 0.01, ***: p < 0.001, ****: p < 0.0001.

To further examine the interaction mode between CODANIN-1 and ASF1A, we focused on the BD-N and HMH-C domains, as only these domains are sufficient to bind to ASF1A. We introduced R195A and R196A mutations (BD-N_mut_) into GST-BD-N, which are predicted to disrupt the two salt bridge interactions observed in our structure (Fig. 2b). GST pulldown experiments revealed that this BD-N mutant completely lost the ability to interact with ASF1A (Fig. 3b). For HMH-C, we generated individual R541A (HMH-C_R541A) and R548A (HMH-C_R548A) mutations in GST-HMH-C, which are predicted to disrupt the salt bridge interaction with the D88 or D54 residues in ASF1A (Fig. 2e). In addition, we generated an L545A/L549A (HMH-C_LL/AA) double mutant to eliminate hydrophobic interactions between HMH-C and ASF1A (Fig. 2e). A GST pulldown assay revealed that these HMH mutations abrogated the interaction with ASF1A, further confirming that CODANIN-1 uses a histone-mimicking mechanism to interact with ASF1 (Fig. 3b). These results revealed that CODANIN-1 utilizes HMH and the B-domain to interact with ASF1A, with BD-N and HMH-C playing major roles.

To quantitatively examine the contribution of each domain (BD-N, HMH-N, HMH-C and BD-C) to ASF1 binding in the context of full-length CODANIN-1, we generated a series of full-length CODANIN-1 mutants and analyzed their binding to ASF1A via biolayer interferometry (BLI) by immobilizing ASF1A as a ligand on the chip surface (Fig. 3c).

First, to examine the contribution of each domain to ASF1A binding, we generated full-length CODANIN-1 mutants in individual domains: BD-N_mut__R195A/R196A (Mut1), HMH-N_mut__L254R (Mut2), HMH-C_mut__L545A/L549A (Mut3) and BD-C_mut__R825A/K826A (Mut4) (Fig. 3c). BLI analysis revealed that wild-type (WT) CODANIN-1 has 12.7×10^4^ k_on_ (M^−1^ s^−1^) for ASF1A binding, with no measurable dissociation (k_off_) (Fig. 3c). Among the single-domain mutants, Mut1 presented a 10-fold reduction in k_on_ (1.95 ×10^4^ M^−1^ s^−1^), whereas the other single-domain mutants (Mut2, Mut3, and Mut4) presented a 2–3-fold decrease in k_on_ (Fig. 3c). However, all the mutants had no measurable dissociation. These data are also consistent with our GST pulldown experiments, which revealed that BD-N is the strongest ASF1A-binding domain (Fig. 3a). Moreover, the BLI data indicate that CODANIN-1 synergistically interacts with ASF1A via multiple domains.

Second, we abrogated the distal or proximal ASF1A binding modules by mutating BD-N together with HMH-N (Mut5: BD-N_mut__R195A/R196A and HMH-N_mut__L254R) and HMH-C with BD-C (Mut6: HMH-C_mut__L545A/L549A and BD-C_mut__R825A/K826A) and examined their binding to ASF1A (Fig. 3c). Mut5 is predicted to lose the interaction with _dist_ASF1A, whereas Mut6 should disrupt binding with _prox_ASF-1 (Fig. 2a). Compared with wild-type CODANIN-1, Mut5 resulted in a 6-fold decrease in k_on_ (2.29×10^4^ M^−1^ s^−1^), and bound ASF1A started dissociating at 3.16×10^−4^ k_off_ (s^−1^), leading to significantly lower binding. Interestingly, Mut6 exhibited similar or slightly greater binding than the wild-type CODANIN-1, with 1.95×10^4^ k_on_ (M^−1^ s^−1^) and 1.18×10^−4^ k_off_ (s^−1^). These data suggest that the distal ASF1A binding module binds more tightly to ASF1A than the proximal ASF1A binding module does and that the distal and proximal ASF1A binding modules independently function in ASF1A binding.

Third, as GST pulldown experiments with individual domain mutants suggested that BD-N and HMH-C are central to the distal and proximal ASF1A binding modules, respectively (Fig. 3a), we generated a double CODANIN-1 mutant (Mut7: BD-N_mut_ and HMH-C_mut_) that disabled BD-N (R195A/R196A) and HMH-C (L545A/L549A). This mutant completely lost ASF1A binding (Fig. 3c), verifying the major roles of BD-N and HMH-C in proximal and distal ASF1A binding, respectively.

Fourth, we examined the collective contribution of the two HMHs to ASF1A binding compared with the two B-domains. Mutations of both BD-N and BD-C (Mut8) strongly abrogated binding to ASF1A (Fig. 3c). However, a mutant (Mut9) with both HMH-N and HMH-C disabled maintained ASF1A binding similar to that of the wild type (Fig. 3c). These results suggest that the B-domains contribute more critically to ASF1A binding than the HMH domains do. While we were performing the BLI experiments, we noticed that some of the mutants (Mut3, Mut4, Mut6, and Mut9) presented a greater response unit (the y-axis in the BLI sensorgram) than did the wild type in BLI (Fig. 3c). Interestingly, all these mutants have mutations in the proximal ASF1A binding module. As BLI also measures the conformational changes in the molecules, we speculate that the loss of interaction between _prox_ASF1A and the proximal ASF1A binding module of CODANIN-1 would release the distal ASF1A binding module bound with _dist_ASF1A, as the distal ASF1A binding module is located in the middle of a large loop and _dist_ASF1A stacks on _prox_ASF1A. This conformational change may affect the BLI measurement. However, we do not have a clear explanation for this observation. Regardless, we note that the results of the GST pulldown experiments with wild-type CODANIN-1 and the full panel of mutants corroborated the results of the BLI measurements (Supplementary Fig. 7).

To further validate the interaction between CODANIN-1 and ASF1A, we designed complementary mutations in ASF1A to disrupt the interaction with CODANIN-1: ASF1A_HMH (D88A, D54A, or V94R) for HMH, ASF1A_BD (ED/AA:E36A/D37A) for the B-domain and ASF1A_HMH+BD (D88A+ED/AA, D54A+ED/AA and V94R+ED/AA) for both the B-domain and HMH interactions. We then performed a GST pulldown assay using ASF1A mutants as the bait and full-length wild-type CODANIN-1 as the prey. Consistent with the BLI results, mutations in the HMH or B-domain binding sites of ASF1A had a modest effect on binding, whereas the double mutation significantly abrogated the binding of ASF1A to CODANIN-1 (Fig. 3d). Together with our cryo-EM structure, these data suggest that CODANIN-1 utilizes its two modules of HMHs and B domains for ASF1A binding and enveloping ASF1A with each domain synergistically.

### CODANIN-1 sequesters ASF1A to prevent histone H3–H4 binding

We revealed that CODANIN-1 has two ASF1A binding modules, each composed of a B domain and an HMH. The HMHs mimic histone H3 to interact with ASF1 and would thus bind ASF1A in its histone-free state, whereas the B-domains could, in principle, bind an ASF1A/H3–H4 complex as well as free ASF1A. The histone-mimicking strategy of CODANIN-1 implies a competitive relationship with histone H3–H4 for ASF1A binding. To test this hypothesis, we carried out pulldown assays with GST-ASF1A in the presence of histone H3–H4 and increasing amounts of CODANIN-1. These assays revealed that histone binding to ASF1A decreased as CODANIN-1 binding increased (Fig. 4a and Supplementary Fig. 8a). These findings indicate that CODANIN-1 sequesters ASF1A and prevents the formation of a complex with histones H3–H4. We also tested this competitive relationship in the presence of CDIN1, but CDIN1 did not significantly affect CODANIN-1 sequestration of ASF1A (Supplementary Fig. 8b).

**Fig. 4.**
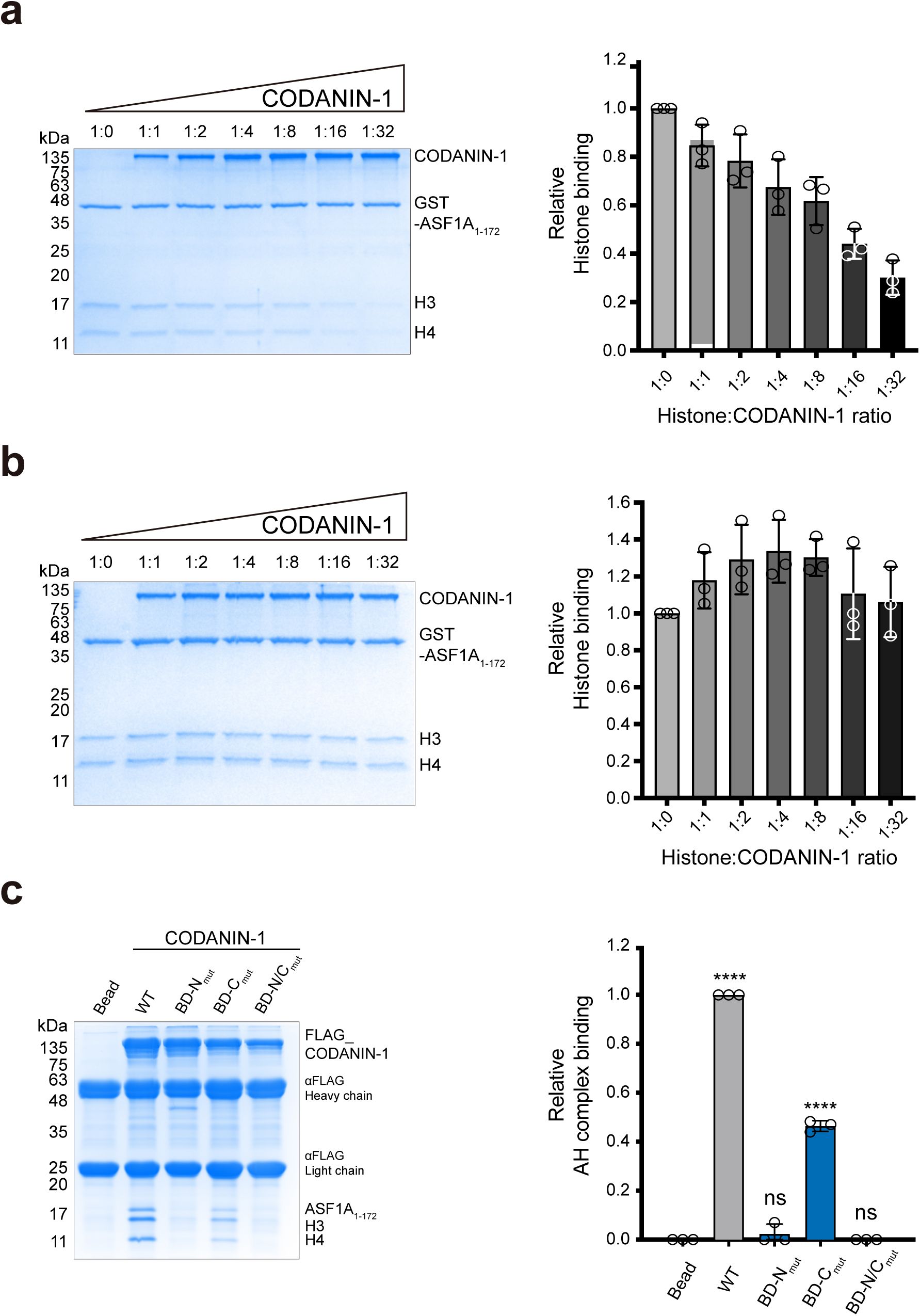
CODANIN-1 sequesters free ASF1A to prevent complex formation with histone H3–H4. **a,** GST-ASF1A pulldown experiment using CODANIN-1 and histone H3–H4 (left panel). The ratio of histone H3–H4 to CODANIN-1 is indicated above each gel lane. Quantification of histone H3–H4 relative to GST-ASF1A (right panel). The value was normalized to the histone H3–H4 band in the absence of CODANIN-1 (1:0 ratio). **b,** Pulldown of CODANIN-1 with a preformed GST-ASF1A/H3–H4 complex (left panel). The ratio of histone H3–H4 to CODANIN-1 is indicated. Quantification of histone H3–H4 relative to GST-ASF1A (right panel). The value was normalized to the histone H3–H4 band in the absence of CODANIN-1 (1:0 ratio). **c**, FLAG-pulldown experiments in which FLAG-tagged full-length CODANIN-1 wild type and mutants (BD-N_mut_, BD-C_mut_, and BD-N/C_mut_) were used for ASF1A/H3–H4 binding. A representative SDS‒PAGE gel is shown in the left panel, and the quantification of the binding (n=3) is shown in the bar graphs. Each dot represents a biological replicate (n=3), and bars and error bars represent the mean±SD. P-values were calculated by one-way analysis of variance (ANOVA) with Dunnett’s multiple comparisons test against ‘Bead’ control (ns: not significant, ****: p < 0.0001, WT: p<0.0001, BD-N_mut_: p=0.4762, BD-C_mut_: p<0.0001, BD-N/C_mut_: p>0.9999).

Next, we tested whether CODANIN-1 can extract ASF1A from a preformed ASF1A/H3–H4 complex. We preincubated GST-ASF1A with H3–H4 and performed pulldown experiments using the GST-ASF1A/H3–H4 complex with increasing amounts of CODANIN-1 as prey (Fig. 4b and Supplementary Fig. 8c). In this setting, while CODANIN-1 still bound to ASF1A/H3–H4, the binding of histone H3–H4 to ASF1A was unaffected by CODANIN-1 (Fig. 4b). This finding suggests that CODANIN-1 cannot deprive ASF1A of its histone H3–H4 dimer, but it can bind to ASF1A/H3–H4.

To investigate whether CODANIN-1 binds to the ASF1A/H3–H4 complex via its B-domains, we mutated the B-domains (BD-N and BD-C), alone and together, in full-length CODANIN-1 (BD-N_mut_, BD-C_mut_, BD-N/C_mut_) and examined binding via FLAG-pulldown experiments (Fig. 4c and Supplementary Fig. 8d). BD-N/C_mut_ completely abrogated the binding to ASF1A/H3–H4 (Fig. 4c), indicating that the B-domains are required for this binding module. In addition, while BD-N_mut_ showed little binding, BD-C_mut_ still bound to ASF1A/H3–H4. This observation is consistent with the finding that GST-BD-N, but not GST-BD-C, is sufficient to bind to ASF1A (Fig. 3a). We further confirmed that CODANIN-1 can bind to ASF1A by the B domain alone using a GST-ASF1A_V94R_ mutant in which the histone-binding interface is disrupted. GST pulldown experiments revealed that both BD-N_mut_ and BD-N/C_mut_ (but not BD-C_mut_) did not bind to ASF1A_V94R_, indicating that BD-N could bind to ASF1A in the absence of the other binding domain (Supplementary Fig. 8e). These data imply that CODANIN-1 has the ability to sequester not only free ASF1A but also histone-bound ASF1A.

### CODANIN-1 dimerization is mediated by the second helical lobe and the C-terminal region

The cryo-EM structure revealed that CODANIN-1 forms a homodimer with a large interface (1,844 Å^2^), which is mediated by LOBE2 (Fig. 1e and Supplementary Fig. 5). Two LOBE2s form a dimer in a saddle joint shape with hydrophobic and electrostatic interactions (Supplementary Fig. 9a). Specifically, at the top of the saddle, the F581 residue from each CODANIN-1 is inserted into a hydrophobic pocket formed by F581, M588, V643, L646, and P647 from the other CODANIN-1 LOBE2 (Fig. 5a and Supplementary Fig. 9a). In addition, R447 and E652 from each molecule form salt bridges (Fig. 5a). To examine the dimerization interface, we introduced F581E and E652A mutations to disrupt the hydrophobic interaction and the salt bridges and analyzed this mutant by size exclusion chromatography (in the absence of CDIN1). However, unexpectedly, this mutant still formed a dimer (Supplementary Fig. 9b). This observation led us to hypothesize that there might be an additional dimerizing domain in the region not observed in the cryo-EM structure and that CODANIN-1 forms a dimer without CDIN1. In our structure, the C-terminal regions of CODANIN-1 (834-1,227 a.a.) and CDIN1 are completely disordered. As part of the C-terminal region (1,000-1,227 a.a.) was previously shown to interact with CDIN1^30–32^, we envisioned that the C-terminal region of CODANIN-1 is involved in CODANIN-1 dimerization in addition to LOBE2. To examine this possibility, we used AlphaFold 3^39,40^ to obtain a predicted model of the C-terminal region (841-1,204 a.a.) of CODANIN-1 in complex with full-length CDIN1 (Supplementary Fig. 10a). The model predicted that the C-terminal region of CODANIN-1 forms two domains. A region including 841-1,000 a.a. formed a coiled-coil structure, whereas a second region of 1,025-1,204 a.a. formed a helical bundle composed of eight α-helices. These two domains are connected by a loop (1,001-1,024 a.a.). The eight helical bundle domains were predicted to form a complex with CDIN1 (Supplementary Fig. 10b). As the domain of a.a. 841-1,000 was predicted to have a coiled-coil structure, we explored the possibility that it could form a homodimer. We performed AlphaFold 3 predictions to generate a dimeric model of the C-terminal region of CODANIN-1 (841-1,204 a.a.) in complex with CDIN1. This dimer model revealed that the two F868 residues from each CODANIN-1 form a ρ-ρ stacking interaction with their aromatic rings, and the R864 and E871 residues from one molecule form salt bridges with E871 and R864 from the other molecules, respectively (Supplementary Fig. 10c). Interestingly, F868 was found to be mutated to I868 in CDA-I disease^33^. However, it is not clear whether the F868I mutation affects dimerization in-vivo. Therefore, on the basis of the predicted model, we generated a dimerization mutant (DM_mut_) with mutations (F868I/E871R) in the C-terminal coiled-coil and mutations (F581E/E652A) in the dimerization interface of LOBE2 in the absence of CDIN1. While mutation in the dimer interface of LOBE2 or the C-terminal coiled-coil region alone did not affect the dimerization of CODANIN-1, a significant portion of the DM_mut_ became monomeric (Fig. 5b and Supplementary Figs. 9b and 9c). Furthermore, negative-stain EM analysis revealed a smaller size of DM_mut_ than of wild-type CODANIN-1 (Fig. 5b). These data suggest that CODANIN-1 has at least two dimerization domains: LOBE2 and the C-terminal region.

**Fig. 5.**
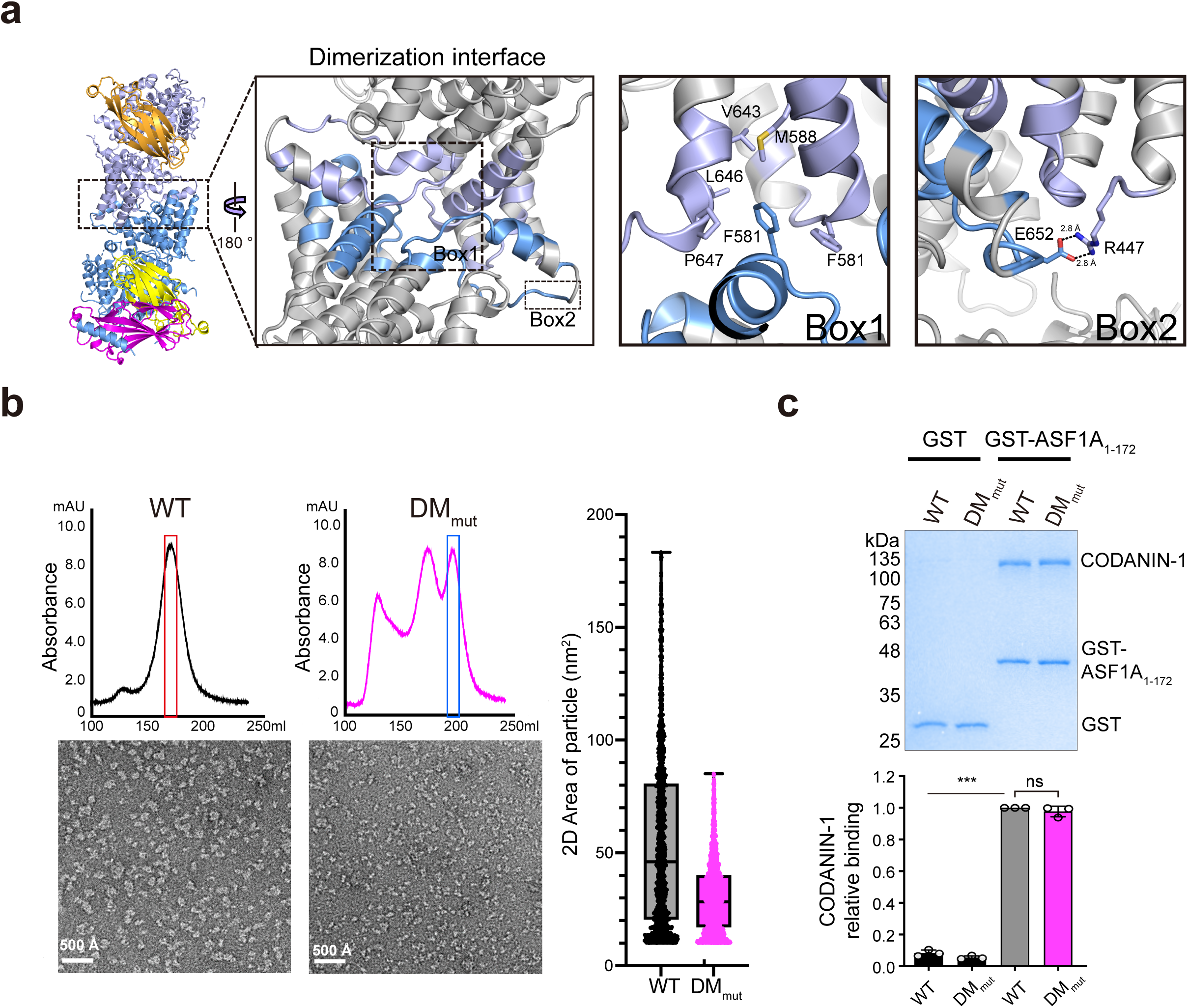
Dimerization of the CODANIN-1 complex. **a,** Detailed view of the dimer interface of the CODANIN-1 complex. The dashed-line boxes (Boxes 1 and 2) indicate the dimerization interfaces of CODANIN-1 and the enlarged view of the CODANIN-1 dimerization interface. The F581 residue of each CODANIN-1 is inserted into a hydrophobic pocket formed by the F581, M588, V643, L646, and P647 residues from the other CODANIN-1 (Box 1). The R447 and E652 residues from the two different CODANIN-1 molecules form a salt bridge (Box 2). **b,** Size-exclusion chromatograms and negative-stain EM micrograph images of wild-type CODANIN-1 and DM_mut_ (left panel). Box and whisker plot with all points shows the 2-dimensional area of the particles (minimum whisker: minimum value, lower quartile: 25% percentile, center line: median, upper quartile: 75% percentile, maximum whisker: maximum value). The number of particles used in the analysis 1405 for WT and 1862 for DM_mut_ (right panel). **c,** GST-pulldown experiment with GST-ASF1A and CODANIN-1 wild-type and DM_mut_ (top panel). The bar graph represents the quantification of CODANIN-1 relative to GST-ASF1A normalized to wild-type CODANIN-1 (right panel). For the bar graphs, each dot represents a biological replicate (n=3), and bars and error bars represent the mean±SD. The statistical P-values were calculated with one-way analysis of variance (ANOVA) with Dunnett’s multiple comparisons test against GST-ASF1A bound WT. (ns: not significant, ** p<0.01, ***: p < 0.001, GST bound WT: p=0.0009, GST bound DM_mut_: p=0.0003, GST-ASF1A bound DM_mut_: p=0.4153).

### The dimerization of CODANIN-1 is not required for ASF1 binding

Our cryo-EM structure with the AlphaFold 3 predicted model indicates that CODANIN-1 forms a dimer through multiple dimerization domains with large interfaces. In addition, CODANIN-1 exhibited dimer formation in gel-filtration chromatography even in the absence of CDIN1. These observations suggest that CODANIN-1 functions as an obligatory dimer with dimerization interfaces located at the ends opposite those of the ASF1 binding domains. However, it is unclear whether the dimerization of CODANIN-1 is critical for ASF1A binding. Therefore, we collected a fraction containing monomeric CODANIN-1 (DM_mut_), confirmed the monomer state by negative stain EM and analytical gel filtration chromatography (Fig. 5b and Supplementary Fig. 9c) and tested the binding of GST-ASF1A to monomeric CODANIN-1 (Fig. 5c). In the GST pulldown experiments, monomeric CODANIN-1 showed comparable binding to that of wild-type dimeric CODANIN-1 toward ASF1A. These data imply that the dimerization of CODANIN-1 is not required for ASF1 binding in-vitro.

### Binding via the HMH and the B-domain is required for CODANIN-1-mediated sequestration of ASF1A in the cytoplasm

Our cryo-EM structure and biochemical analysis revealed the interaction mode between CODANIN-1 and ASF1A. Next, we aimed to explore the interaction in cells and, more specifically, the ability of CODANIN-1 to sequester ASF1A in the cytoplasm and thereby control the flow of newly synthesized histones to chromatin. We utilized the Flp-In system to integrate inducible siRNA-resistant CDAN1 constructs into U-2-OS cells as a complementation system^29^ (Supplementary Fig. 11a). As CODANIN-1 is expressed at low levels in cells, we tagged the mutant constructs with a destabilization domain to keep the mutant protein levels low^41^. However, even with this approach, the complemented proteins were more highly expressed than the endogenous wild-type CODANIN-1 was. The endogenous protein was depleted by siRNA treatment to address the functionality of CODANIN-1 mutants in the absence of endogenous protein. We validated the knockdown of endogenous CODANIN-1 (Fig. 6a), with siRNA #2 having the strongest effect and therefore being used for the complementation experiments.

**Fig. 6.**
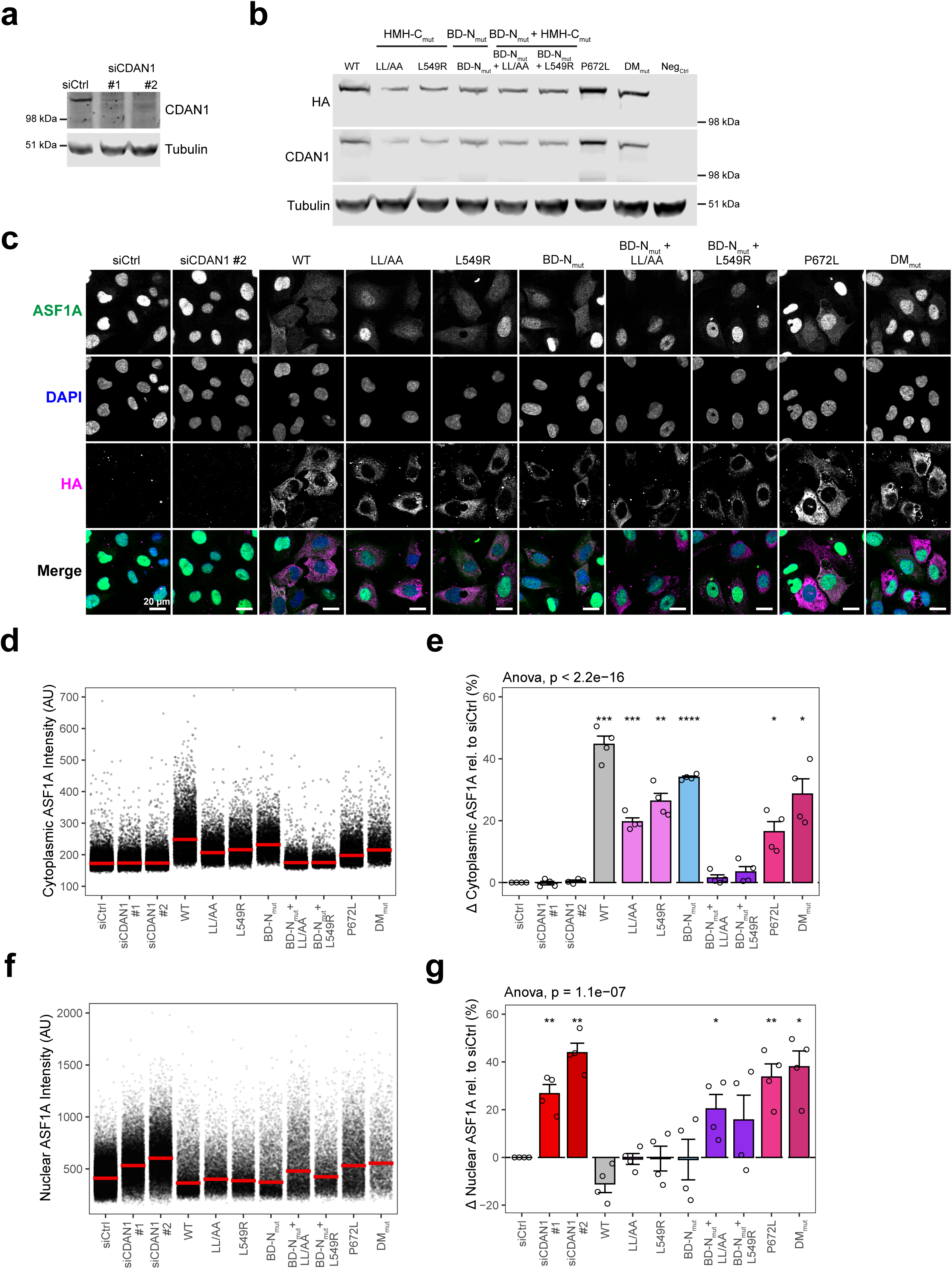
One B-domain and one HMH are required for cytoplasmic sequestration of ASF1A is required for cytoplasmic sequestration by CODANIN-1. **a**, Western blot of endogenous CODANIN-1 after 48 hours of siRNA treatment with control siRNA (siCtrl) or siRNA targeting CDAN1 (siCDAN1 #1 and #2). **b**, Western blot showing the expression of HA-tagged CODANIN-1 after 24 hours of treatment with doxycycline and Shield1. WT: wild type, B_mut_: B domain mutant, DM_mut_: dimerisation mutant. The Western blot is representative of n=2 biological replicates. **c**, Representative images of ASF1A localization in cell lines complemented with exogenous WT and mutant CODANIN-1 as indicated. The siCtrl and siCDAN1 #2 panels represent the parental cell lines with control or CDAN1 knockdown, respectively. Each cell line was treated with CDAN1#2 siRNA for 48 hours, and doxycycline and Shield1 were added for the last 24 hours to induce the expression of exogenous CODANIN-1. **d**, Cytoplasmic ASF1A intensity quantified in individual cells from a single biological replicate as in c. For complementation with exogenous CODANIN-1, only HA-expressing cells were included. The bars repserent the median and interquartile range of each condition. **e**, Relative change in cytoplasmic ASF1A intensity compared with that of the control. For each condition the difference in median cytoplasmic ASF1A intensity compared with the median of siCtrl was calculated (% change). Each dot represents a biological replicate (n=4), and the bars and error bars represent the means±SDs. P-values were calculated by two-sided paired t-test against siCtrl (* p<0.05, ** p<0.01, *** p<0.001, **** p<0.0001, WT: p=4.4×10^−4^, LL/AA: p=5×10^−4^, L549R: p=1.6×10^−3^, B_mut_: p=4.8×10^−5^, P672L: p=1.5×10^−2^, DM_mut_: p=1.2×10^−2^). **f,g**, Nuclear ASF1A intensity quantified in HA-expressing cells using the same methods as in d,e, A minimum of 2000 cells HA-expressing were considered per condition. Each dot represents a biological replicate (n=4), and bars and error bars represent the mean±SD. P-values were calculated by two-sided paired t-test against siCtrl (* p<0.05, ** p<0.01, *** p<0.001, **** p<0.0001, siCDAN1 #1: p=8.3×10^−3^, siCDAN1 #2: p=1.5×10^−3^, B_mut_+LL/AA: p=4×10^−2^, P672L: p=8.1×10^−3^, DM_mut_: p=1×10^−2^).

Our cryo-EM and biochemical studies revealed that the BD-N and HMH-C domains are the major binding sites for the distal and proximal ASF1 binding modules, respectively, and mutation of these domains completely abrogated CODANIN-1 binding to ASF1A in-vitro. Therefore, we decided to focus our cellular assays on BD-N and HMH-C domain mutants. To assess the ability of CODANIN-1 to sequester ASF1A in the cytoplasm^29^, we imaged the localization of ASF1A in response to the knockdown of endogenous CODANIN-1 and complementation with wild-type or mutant CODANIN-1 (Fig. 6b). We used HA-tagged CODANIN-1 staining to identify cells with efficient complementation and validated comparable expression levels across the cell lines (Supplementary Fig. 11b, c). Cells expressing wild-type CODANIN-1 presented a shift in ASF1A to the cytoplasm (Fig. 6c-e) and depletion of ASF1A from the nucleus (Fig. 6f, g), in agreement with the ability of CODANIN-1 to sequester ASF1 in the cytoplasm upon overexpression^29^. Single mutants of HMH-C (HMH-C_LL/AA: L545A/L549A; HMH-C_L549R) or BD-N (BD-N_mut_: R195A/R196A) presented a cytoplasmic ASF1A signal but at a slightly lower level than that of wild-type CODANIN-1. These data are consistent with our biochemical data showing that a single mutation in HMH-C or BD-N did not completely abrogate ASF1A binding.

Notably, when the double mutants (BD-N_mut_+HMH-C_LL/AA and BD-N_mut_+HMH-C_L549R) were expressed, ASF1A remained entirely nuclear (Fig. 6c-e), which is consistent with our BLI experiments showing that Mut7 (BD-N and HMH-C mutant) completely lost the ability to bind to ASF1A (Fig. 3c). These data demonstrate that the binding mode of CODANIN-1 via both the B-domain and HMH is critical for ASF1A interaction and sequestration in cells. We also examined the effect of CODANIN-1 dimerization on ASF1A binding. DM_mut_ impaired sequestration of ASF1A (Figs. 6d and 6e), indicating that failure to dimerize CODANIN-1 impairs its ability to bind and sequester ASF1A in a cellular context, despite the ability of monomeric CODANIN-1 to interact with ASF1A in-vitro. This was not related to the localization of DM_mut_, as this was comparable to that of wild-type CODANIN-1 (Supplementary Figs. 11b, c). The CDA-I disease mutation P672L is located in the dimerization interface, although the mutant still forms a dimer in-vitro (Supplementary Fig. 9b). Therefore, we also examined the ability of the pathogenic P672L mutant to sequester ASF1A (Fig. 6d and 6e). This mutant presented reduced ASF1A sequestration, similar to that of DM_mut_ (Figs. 6d and 6e). In summary, we demonstrate that the dual interaction mode of CODANIN-1 via HMH-C and BD-N is required for ASF1A binding in cells and its ability to sequester ASF1A in the cytoplasm. We further showed that CODANIN-1 dimerization also influenced this process.

## Discussion

CODANIN-1 is unique in its role as a negative regulator of histone supply^29,42^, but the mechanism by which CODANIN-1 binds and sequesters ASF1 remains unknown. Here, we report the atomic resolution cryo-EM structure of the CODANIN-1_ASF1A complex. We found that CODANIN-1 forms a dimer in which each monomer interacts with two molecules of ASF1A through proximal and distal binding modules. This finding is consistent with a recent report on the structure of the CODANIN-1_ASF1A complex^43^. The ability of the CODANIN-1 complex to bind multiple ASF1 molecules highlights its capacity as a negative regulator of ASF1A histone import activity. We demonstrated that CODANIN-1 binds ASF1A in a competitive manner with histones H3–H4 but can also bind a preformed ASF1A/H3–H4 complex, suggesting two possible modes for CODANIN-1 regulation of ASF1A activity. We further showed that the combined proximal and distal binding of ASF1A by CODANIN-1 is essential for the cytoplasmic sequestration of ASF1A in cells, demonstrating the importance of this interaction in regulating the histone supply. This finding strengthens our understanding of histone supply regulation and reveals the mechanisms underlying CODANIN-1-mediated regulation of ASF1A activity.

The structure of the CODANIN-1_ASF1A complex revealed that CODANIN-1 has two ASF1A binding modules composed of two HMH helices (HMH-N and HMH-C) and two B-domains (BD-N and BD-C). These domains are highly conserved among species (Supplementary Fig. 4), implying conservation and functional importance of ASF1A regulation by CODANIN-1. The dual-binding mode by the HMH and the B-domain occludes both major interaction sites of ASF1A, reminiscent of early work suggesting a similar dual-binding mechanism mode of Rad53 interaction with yeast Asf1^44^, which would be interesting to revisit in light of our findings. CODANIN-1 binding prevents the interaction of ASF1A with histones via HMH and with other chaperones by the B domain, thereby sequestering ASF1A in a fully inactive state.

Consistent with the importance of multiple binding modes for ASF1A inactivation, abrogating both the distal and proximal ASF1A binding modules by mutating BD-N and HMH-C was required to fully prevent ASF1A sequestration (Fig. 6e). Our BLI data revealed that CODANIN-1 binds with 12.7×10^4^ k_on_ (M^−1^s^−1^) with no detectable dissociation (k_off_), indicating very strong binding of ASF1A (Fig. 3c). Considering that ASF1A sequestration should be dynamic to regulate the histone supply in response to stimuli, it would be interesting to examine whether there is a regulatory mechanism to release ASF1A from CODANIN-1 that has yet to be identified. The fact that CODANIN-1 has HMH occupying the histone-binding surface of ASF1A implies that CODANIN-1 prevents ASF1A from forming a histone-bound complex. It would also prevent the binding of TLK2, which binds to ASF1 through a similar mechanism using a histone-mimicking motif^45^ and acts as a positive regulator of the histone supply. Therefore, CODANIN-1 HMH binding would strongly inhibit the ASF1-mediated histone supply, preventing both activating phosphorylation by TLK2 and histone binding. DNAJC9 is suggested to support ASF1 in the histone supply chain^12^, and investigating how ASF1 inactivation by CODANIN-1 affects the activities of DNAJC9 and other chaperones in the histone supply chain would be interesting.

The B-domain is also found in other ASF1A-binding proteins, such as CAF-1 and HIRA^27,28,44,46,47^, and the binding mode of the CODANIN-1 B-domains to ASF1A is similar to that of these downstream chaperones. Interestingly, in our cryo-EM structure, BD-N (a.a. 193-202) and HMH-N (a.a. 249-260) are the only structured sections inside a large disordered region (a.a. 69-284), suggesting that this region likely becomes ordered upon interaction with ASF1A. Therefore, without ASF1A binding, the entire region (a.a. 69-284) containing the BD-N and the HMH-N would be flexible. In contrast to BD-N, BD-C (a.a. 820-833) is the last section of a structured region, and the entire C-terminal region beyond BD-C is disordered (Fig. 1a). It is plausible that this large loop embedding BD-N and the HMH-N may capture free ASF1A to be tethered to CODANIN-1.

The ability of CODANIN-1 to also bind the ASF1A-histone complex (Figs. 4b and 4c) suggests that CODANIN-1, via its B-domain, could be a mechanism to regulate ASF1A delivery of histones to downstream chaperones. This could not only be used to sequester histone-bound ASF1A outside of the nucleus, as occurs when CODANIN-1 is overexpressed (Fig. 6c), but may also be a function of the endogenous nuclear pool of CODANIN-1, which predominantly localizes to the nucleolus^48^. CODANIN-1 binding to ASF1A/H3–H4 inhibits the handover of histones to downstream B-domain-containing chaperones such as CAF-1 and HIRA. This could lead to sequestration of ASF1A/H3–H4 in the nucleus and provide a more dynamic mechanism for regulating the histone supply by acting downstream of histone import. This may be especially important in conditions where the histone supply must be strongly inhibited, for example, during erythrocyte differentiation and enucleation, where excess delivery of acetylated histones might counteract the histone deacetylation^49,50^ and removal^51,52^ critical for the process. The identification of the CODANIN-1_ASF1 binding mode now opens an avenue for structure‒function studies to address how this inhibitory axis feeds into the functions of chaperones such as CAF-1, HIRA, and DNAJC9 in different cell types.

In the cryo-EM structure, a large portion of the C-terminal region of CODANIN-1 and CDIN1 bound to this region is not visible, suggesting that this part of the complex is dynamic. Interestingly, our structural and biochemical analysis of the C-terminal region revealed a role for CODANIN-1 dimerization. Mutations in CODANIN-1 and CDIN1 cause CDA-I disease, suggesting that they act in the same functional axis^33,34,48,53^. However, CODANIN-1 alone was sufficient to bind ASF1 in our in-vitro binding assay. The contribution of CDIN1 to CODANIN-1 function in the regulation of histone supply is thus unclear, and further structural and mechanistic studies are needed to elucidate CDIN1 function. We identified that the positions of pathogenic mutations involved in CDA-I are clustered into 6 regions^48^. Interestingly, three regions are located toward the C-terminus in the large dimerization domain of CODANIN-1 and in CDIN1^48^. CODANIN-1 interacts with CDIN1 through its C-terminal domain (1,001-1,227 a.a.), which contains the CDA-I disease mutant (R1042W) with impaired capacity for ASF1A binding^29^. The CDA-I mutation (P672L) and the DM_mut_ assessed in this study resulted in a similar reduction in the ability to sequester ASF1A (Figs. 6d and 6e). Although DM_mut_ retained ASF1A-binding ability in-vitro (Fig. 5c), the reduced sequestration in-vivo could indicate that the formation of a larger CODANIN-1 complex plays an important role in ASF1A regulation in a cellular context. The inability to efficiently form a larger CODANIN-1 dimer complex could affect the recruitment of other factors, leading to untimely ASF1 release, or might alter the association of CDIN1 with the complex. The prevalence of disease-causing mutations in the dimerization and CDIN1 interaction interfaces, along with the reduction in ASF1A sequestration in a pathogenic CODANIN-1 mutant (this study and Ask et al.)^29^, suggest that some CDA-I disease-causing mutants affect the integrity of the loss of CODANIN-1_CDIN1 complex integrity, which contributes to CDA-I development potentially in part through misregulation of ASF1A. The spongy heterochromatin and nuclear abnormalities of CDA-I^35,37,54^ may also suggest that defective chromatin regulation may underlie the disease. This could be due to an inability to sequester ASF1A outside the nucleus or by altering the interactions of ASF1A with other chaperones within the nucleus. CODANIN-1_CDIN1 could also have further unexplored activities that could contribute to disease pathogenesis; for example, DNA binding or nuclease activities were first suggested for CDIN1 on the basis of sequence similarity^34^. Further exploration of the structure‒function of the CODANIN-1_CDIN1 complex and its roles in chromatin regulation during hematopoiesis will be important for understanding and treating CDA-I.

## Methods

### Purification of CODANIN-1 complex

CODANIN-1 and CDIN1 were cloned into a pFastBac1 vector with a C-terminal 6xHis tag and a TEV protease cleavage site (ENLYFQG) before the His tag for CODANIN-1, and with no tag for CDIN1. The cloned plasmids were transformed into DH10Bac competent cells to generate bacmid DNAs. Bacmid DNAs were transfected into *Spodoptera frugiperda* 9 (Sf9) cells to produce baculoviruses harboring CODANIN-1 and CDIN1. These baculoviruses were amplified to passage 3 according to the Bac-to-Bac Baculovirus Expression Kit (Thermo Fisher Scientific). The two baculoviruses were co-infected into Sf9 cells and harvested 48 hours post-infection in a lysis buffer (50 mM Tris, pH 8.0, 500 mM NaCl, 5% glycerol) containing a protease inhibitor cocktail (Roche).The Sf9 cells were lysed by repeated freezing and thawing cycles and cell debris were cleared with a centrifugation at 27,216 g for 2 hrs. ASF1a_1-172_ was cloned into pET21a vector with no tag and was expressed from *Escherichia coli (E. coli)* BL21 Codon Plus(DE3)-RILP. The bacteria cells were harvested in a lysis buffer and lyzed by sonication and the debris were cleared with a centrifugation at 27,216 g for 2 hours. The prepared Sf9 and bacteria cell lysates were mixed and the mixture was loaded into Ni-NTA resin (Qiagen) packed gravity-flow column. The column was washed with a washing buffer (50 mM Tris pH 8.0, 500 mM NaCl, 5% glycerol, 20 mM Imidazole) and the bound protein was eluted with an elution buffer (50 mM Tris pH 8.0, 500 mM NaCl, 5% glycerol, 150 mM Imidazole). The eluted sample was then diluted up with the four volume of the sample with 50 mM Tris HCl pH8.0 and loaded into HiTrap Q HP (Cytiva). The protein was eluted with a salt gradient from 0 to 1 M NaCl. The fraction containing CODANIN-1 complex was further purified with a size exclusion chromatography (HiLoad 16/600 Superose 6 pg (Cytiva)) equilbriated with a buffer (50 mM Tris pH8.0, 500 mM NaCl, 5% glycerol). The fractions containing CODANIN-1_ASF1A_CDIN1 fractions were collected and concentrated by Amicon Ultra Centrifugal Filter 100 kDa MWCO (Merck Millipore) up to 3mg/mL. The sample was stored at −80°C after flash-freezing into liquid nitrogen. CODANIN-1 mutants were purified similarly.

### Cryo-EM structure determination

To determine the cryo-EM structure of the CODANIN-1 complex, the complex was further separated with a GraFix method^55^. Briefly, the sample was separated by sucrose gradient ultracentrifugation with the gradients of 10 to 30 % sucrose and 0 to 0.2 % glutaraldehyde at 97,311 g at 4°C for 18hrs using an Optima™ XL-100K ultracentrifuge (Beckman Coulter). The fractions containing the complex were pooled and the sucrose was removed by using Amicon Ultra Centrifugal Filter, 100 kDa MWCO, 0.5ml (Merck Millipore). To overcome an orientation bias problem of the sample with conventional Quantifoil grids, the sample was then loaded on a graphene oxide covered (homemade) holey carbon grid (Quantifoil R1.2/1.3, copper, 300 mesh) and plunge-frozen using a Vitrobot Mark IV (FEI/Thermo Fisher Scientific) under 100% humidity at 15°C with −13 blotting force and 3 sec blotting time. The micrographs were collected using a Titan Krios G4 300 keV Transmission Electron Microscope (Thermo Scientific) equipped with a BioQuantum K3 detector (Gatan) at Institute of Membrane Proteins (IMP). Total 4,282 micrographs were collected with parameters of 1.1 apix, 58.9e/Å^2^ total electron dose over 50 frames, 18 eV of energy filter slit width, and defocus range of −0.8 μm to −2.4 μm (0.2 μm/steps). The data were processed with CryoSPARC v4.2.1^56^. The micrographs were preprocessed with Full-Frame Motion Correction and CTFFIND4 embedded in CryoSPARC^57,58^. The total of 4,075 micrographs were selected based on the proper defocus value, the degree of astigmatism and the motion distance and curvature. From the selected micrographs, particles were picked by blob-picking followed by template-based picking and subjected to four rounds of 2D classification. 36,361 good particles were selected and used as a training data set for Topaz picking^59^. Out of the total 1,592,796 particles picked, 524,649 particles were selected for reconstructing initial models and for 3D classification. After iterative heterogeneous refinement followed by local motion correction^60^, a final set of 56,039 particles were used to reconstructing a 4.19 Å resolution cryo-EM map of CODANIN-1 complex with C1 symmetry. After two rounds of global CTF refinement and non-uniform refinement^61^, the final map reached at 3.75 Å resolution calculated by gold-standard Fourier shell correlation (cut-off at 0.143). Map sharpening was automatically operated in CryoSPARC 3D refinement with a calculated B-factor. The local resolution map was generated in CryoSPARC local resolution estimation.

The atomic model was built based on final refinement map. The structure of ASF1A (PDB ID 2IO5) was used as an initial model for ASF1A^26^. This model was manually fitted into the density map using ChimeraX^62^. Subsequently, manual rebuilding with Cα tracing and side chain fitting was performed in Coot^63^ and adjusted through rigid-body refinement in Phenix^64^. This was followed by real-space refinement in Phenix, incorporating minimization global, local grid search, ADP, occupancy refinement with secondary structure restraints and Ramachandran restraints. Iterative adjustments were then made to eliminate Ramachandran outliers, Rotamer outliers and Cβ outliers, and to improve the model geometry by coot and Isolde 1.3^65^. The final statistics (Table 1) were calculated through comprehensive validation in Phenix.

### Protein expression

We generated GST fusion peptides containing either each BD-N (193-202 a.a.), HMH-N (249-260 a.a.), HMH-C (539-550 a.a.), or BD-C (820-833 a.a.). Additionally, we also generated HMH-C mutants (R541A, R448A, LL/AA: L545A/L549A), and a BD-N mutant (B_mut_ : R195A/R196A) using a modified version of the pET22b vector (Novagen). Subsequently, to expressed all of GST-fused HMH and mutants and GST-fused B-domain and mutant, each clone was transformed into E. coli strain BL21 Codon Plus (DE3)-RILP cells. The cells were cultured in a shaking incubator at 37°C and 180rpm. Upon reaching an OD600 of 0.6, IPTG was added to a final concentration of 0.5 mM for induction. Following induction, for all cell batches were cultured for 3 hours and then harvested in lysis buffer (50 mM Tris, pH 8.0, 500 mM NaCl, 5% glycerol, 1 mM PMSF). Sonication was employed to lyse the cells, and subsequent removal of cell debris by centrifugation at 27,216 g for 1 hour. For the prepared bacterial cell lysates, 2mL of Glutathione Sepharose 4B resin (Cytiva) was added and incubated at 4°C for 2 hours on a tube roller mixer. Consequently, four times of washing with wash buffer (50 mM Tris, pH 8.0, 500 mM NaCl, 5% glycerol) followed by the resin loading into gravity-flow column. The column was washed with a washing buffer (50 mM Tris pH 8.0, 500 mM NaCl, 5% glycerol) and the bound protein was eluted with an elution buffer (50 mM Tris pH 8.0, 150 mM NaCl, 5% glycerol, 10 mM Glutathione reduced). The eluted sample was then diluted up with the three volume of the sample with 50 mM Tris HCl pH8.0 and loaded into HiTrap Q HP (Cytiva). The protein was eluted with a salt gradient from 0 to 1 M NaCl. The fractions containing GST fusion HMH or GST fusion B-domain were collected and concentrated by Amicon Ultra Centrifugal Filter 10 kDa MWCO (Merck Millipore) up to 3 mg/mL. The sample was stored at −80°C after flash-freezing into liquid nitrogen.

For the GST fusion ASF1A_1-172_, we cloned truncated ASF1A (1-172 a.a.) into pGEX-4T-1 vector. Following this, we introduced mutations on the HMH and B-domain interaction surface of ASF1A: D88A, D54A, V94R for HMH interaction, E36A/D37A (ED/AA) for B-domain interaction. To expressed GST-ASF1A_1-172_ and its mutants, we transformed these clones into E. coli strain BL21 Codon Plus (DE3)-RILP cells. The protein expression and preparation steps were identical to those used for the aforementioned GST fusion peptides. GST-ASF1A variants were concentrated using Amicon Ultra Centrifugal Filter 30 kDa MWCO (Merck Millipore) to a 2.5 mg/mL. The concentrated proteins were then flash-frozen in liquid nitrogen and stored at −80°C.

### GST pulldown assay

To verify the interaction between HMHs and ASF1A, as well as between the B-domains and ASF1A, we conducted a GST pulldown assay. In this assay, 25 μl of Glutathione Sepharose 4B resin (Cytiva) was mixed with 125 μg of purified GST fusion HMH or B-domain in a total 500 μl volume of incubation buffer (50 mM Tris, pH 8.0, 500 mM NaCl, 5% glycerol). The mixture was incubated at 4°C for 2 hours on a tube roller mixer. The beads were then washed four times with wash buffer (50 mM Tris, pH 8.0, 150 mM NaCl, 5% glycerol, 0.5% Triton X-100), followed by the addition of 150 μg of ASF1A_1-172_ in a total volume of 500 μl of incubation buffer. This mixture was incubated for 2 hours at 4°C on a tube roller mixer. Post-incubation, the beads underwent four washes with wash buffer to eliminate non-specific binding. Following this, SDS sample buffer was added, and SDS-PAGE was performed using a 16% acrylamide gel. The intensity of the ASF1A bands was measured using Image Lab Software (Bio-Rad). The GST pulldown assays for HMH and B-domain mutants were conducted in the same manner.

To verify the interaction between full-length CODANIN-1 and its mutants with ASF1A, we designed a GST pulldown assay. In this assay, 25 μl of Glutathione Sepharose 4B resin (Cytiva) was mixed with purified GST fusion ASF1A and CODANIN-1 to reach a final concentration of 100 nM each in a total 500 μl volume of incubation buffer (50 mM Tris, pH 8.0, 500 mM NaCl, 5% glycerol). The mixture was then incubated at 4°C for 2 hours on a tube roller mixer. The beads were washed four times with wash buffer (50 mM Tris, pH 8.0, 500 mM NaCl, 5% glycerol, 0.5% Triton X-100), and SDS sample buffer was added to denature the proteins. SDS-PAGE was carried out using a 10% acrylamide gel, and the gel intensity was measured using Image Lab Software (Bio-Rad). The GST pulldown assays for mutant ASF1A and CODANIN-1 were conducted in the same manner.

To verify the competitive relationship between CODANIN-1 and histone H3-H4 over the ASF1 binding, we conducted GST pulldown-based competition assay. In this assay, 25μl of Glutathione Sepharose 4B resin (Cytiva) was mixed with a final concentration of 100 nM purified GST fusion ASF1A in a total volume of 500 μl incubation buffer (50 mM Tris, pH 8.0, 500 mM NaCl, 5% glycerol). Subsequently, histone H3-H4 was added to a final 100nM, and CODANIN-1 was added at doubling molar ratios ranging from 0 to 32 times that of histone H3-H4 (0 nM to 3200 nM). The mixture was incubated for 2 hours at 4°C on a tube roller mixer. Following incubation, the resin was washed four times with wash buffer (50 mM Tris, pH 8.0, 500 mM NaCl, 5% glycerol, 0.5% Triton X-100), and SDS sample buffer was added to denature the proteins. SDS-PAGE was carried out using a 10% acrylamide gel, and the gel intensity was measured using Image Lab Software (Bio-Rad).

In the histone depriving assay, we utilized a final 100 nM of the pre-formed GST-ASF1A and histone H3-H4 complex as a bait, instead of free-ASF1A. CODANIN-1 was subsequently added at doubling molar ratios, ranging from 0 nM to 3200 nM (0 to 32 times the concentration of histone H3-H4). The remaining conditions were identical to those described in the aforementioned competition assay.

### FLAG pulldown assay

To verify the interaction between full-length CODANIN-1 and its mutants with ASF1A/H3-H4 binding, a FLAG pulldown assay was designed. In this assay, 25 μl of Anti-DYKDDDDK G1 Affinity Resin (GenScript) was mixed with 50 μg purified FLAG fusion CODANIN-1 in a total 500 μl volume of incubation buffer (50 mM Tris, pH 8.0, 500 mM NaCl, 5% glycerol). The mixture was incubated at 4°C for 2 hours on a tube roller mixer. The beads were washed four times with wash buffer (50 mM Tris, pH 8.0, 500 mM NaCl, 5% glycerol, 0.5% Triton X-100). After washing, 40 μg of purified ASF1A/H3-H4 complex was added in a total volume of 500 μl incubation buffer. The mixture was then incubated at 4°C for 1.5 hours on a tube roller mixer. The beads were washed four times with wash buffer, and SDS sample buffer was added to denature the proteins. SDS-PAGE was carried out using a 12% acrylamide gel, and the gel intensity was measured using Image Lab Software (Bio-Rad).

### Biolayer interferometry (BLI)

We employed purified 6xHis-tagged ASF1A_1-172_ as the ligand and CODANIN-1 variants, including wild type and mutants: Mut1 (BD-N_mut_), Mut2 (HMH-N_mut_), Mut3 (HMH-C_mut_), Mut4 (BD-C_mut_), Mut5 (BD-N_mut_ / HMH-N_mut_), Mut6 (HMH-C_mut_ / BD-C_mut_), Mut7 (BD-N_mut_ / HMH-C_mut_), Mut8 (BD-N_mut_ / BD-C_mut_), and Mut9 (HMH-N_mut_ / HMH-C_mut_) as analytes. Each sample was stored in a working buffer (50 mM Tris, pH 8.0, 500 mM NaCl, 5% glycerol, 1X kinetics buffer Octet Kinetics Buffer (Sartorius)). BLI measurements were conducted using the Octet R8 (sartorius). Initially, a baseline was established for 60 seconds using the working buffer. Following this, 4 μM of 6xHis-tagged ASF1A_1-172_ was immobilized on nickel-charged tris-nitriloacetic acid (Tris-NTA) biosensor (Sartorius) for the 90 seconds. A second baseline was then established for 60 seconds with the working buffer. Subsequently, the CODANIN-1 variants were introduced during a 240-second association step. Finally, the biosensor was immersed in the washing buffer for a 480-second dissociation step. To determine the binding kinetics, CODANIN-1 underwent seven BLI runs with serial dilutions ranging from 512 nM to 8 nM. Binding kinetics were analyzed by the Octet BLI Analysis 12.2.2.4 software package (Sartorius).

### Cell culture

U2OS cells were cultured in Dulbecco’s Modified Eagle Medium (Gibco) supplemented with 10% FBS and 1% penicillin/streptomycin. Cells were passaged every 4-6d when they reached approximately 80% confluency. FlpIn U2OS cells containing the mutant CDAN1 constructs were maintained with 100 μg/ml hygromycin.

### Mutant CDAN1 Plasmid Generation

A plasmid containing siRNA-resistant FLAG-HA-CDAN1 cDNA(Ask et al., 2012) on an FRT/TO backbone for Flippase integration was used as the original template. The Shield-1 destabilization domain was fused to the N-terminus after the FLAG-HA tag by megaprimer cloning and the resulting construct was sequenced to check for mutations. Q5 PCR-based site-directed mutagenesis was then used to introduce each point mutation, where back to back primers were used to amplify the whole plasmid and introduce a mutation. Primers were designed using NEBasechanger and are listed in Supplementary Table 3. PCR was performed according to manufacturer instructions with an annealing temperature of 72C. 1μl of PCR product was treated with 1μl T4 PNK, 1μl T4 PNK buffer (NEB) supplemented with 10 mM ATP and 1μl DpnI (NEB) for 1 hour, before adding 1μl Quick Ligase for 15 min prior to transformation of Top10 bacteria. Colonies were picked and plasmids extracted by MiniPrep kit (Qiagen) and the mutation sites were sequenced to confirm successful integration. Sanger sequencing was performed by Eurofins Genomics TubeSeq service and a list of plasmids generated is available in Supplementary Table 2.

### Transfection and selection

5×10^5^ U2OS cells were seeded in wells of a 6-well plate 24 hours prior to transfection. Medium was replaced with fresh medium prior to transfection. Transfection was performed using Lipofectamine 3000 (ThermoFisher), with 250ng of FRT/TO plasmid and 2 μg of pOG44 containing Flippase. Control wells containing FRT/TO plasmid or pOG44 alone were also transfected, along with an untransfected control. Transfection medium was replaced after 24 hours for full medium. 24 hours later, cells were seeded on a 10cm plate with 200 μg/ml hygromycin, and hygromycin-containing medium was replaced every 3-4d until complete death of the control cells.

### siRNA transfection

siRNA transfections were performed with Lipofectamine RNAimax (ThermoFisher) according to manufacturer instructions. siRNAs were added to a final concentration of 10nM. For western blot, the siRNA mix was added dropwise to the well 48 hours prior to harvesting. For the transfections in 96-well plates for microscopy, the siRNA mix was added to each well before seeding, 48 hours before fixation. Control siRNA (siCtrl) used was MISSION siRNA Universal Negative Control #1 (Sigma, SIC001). The CDAN1-targeting siRNAs were siCdan1 #1 (Sigma): 5′-CGUAGAGUUCGUGGCAGAAAGAAUU-3′ (sense strand), siCdan1 #2: ON-TARGET plus SMART pool (Dharmacon).

### Western Blot

2×10^5^ U2OS cells were seeded 48 hours prior to siRNA transfection in 6-well plates. siRNA treatment was performed 48 hours prior to harvest. In the complementation cell lines, expression of the CDAN1 construct was induced with 100 ng/μl doxycycline and 50nM Shield1 for 24 hours prior to harvesting. Cells were harvested by washing 2x in cold PBS and lysing in 50 μl Laemmli sample buffer (50 mM Tris-Hcl pH 6.8, 4% SDS, 10% glycerol, 0.1% bromophenol blue, and 25 mM DTT) and treated for 1 hour with 1 μl of benzonase and 0.5 μl of 1M MgCl2. 5 μl of protein lysate was separated by gel electrophoresis on a NuPAGE 4-12% Bis-Tris Gel (ThermoFisher) and transferred by semi-dry transfer in NuPAGE transfer buffer (ThermoFisher) supplemented with 20% ethanol. Transfer efficiency was assessed by Ponceau staining. Membranes were blocked in 5% milk in phosphate-buffered saline with 0.01% tween-20 (PBST) for 1 hour at room temperature, before being incubated with antibodies in milk overnight at 4°C. Membranes were washed 3x in PBST before incubating for 1 hour with secondary antibodies at RT. Secondary antibodies used were IRDye 680RD Goat anti-Mouse IgG and IRDye 800CW Goat anti-Rabbit IgG (Licor). Once the secondary antibodies were added, all incubations were performed in the dark. After secondary incubation, the membranes were washed 3x in PBST and imaged using a LiCOR Odyssey M. All western blots were performed with 2 biological replicates.

### Microscopy

7.5 ×10^4^ U2OS cells resuspended in 150 μl of medium were added to each well of a 96 well plate containing 50 μl of siRNA transfection mix. Medium was replaced with new medium containing 100 ng/μl doxycycline and 50nM Shield1 24 hours prior to harvest. Cells were fixed 48 hours after seeding. To fix, the plates were washed in cold PBS and incubated with 4% paraformaldehyde (PFA) for 15 minutes. PFA was removed and the plates washed 3x in PBS. Wells were filled with 100 μl of cold PBS, sealed with parafilm, and kept in the fridge until further processing. Each experiment was performed with 4 biological replicates with cells cultured separately for a minimum of 2 passages. Cells were permeabilized in 0.5% Triton-X for 15 minutes. All incubation steps were performed with gentle shaking. Each well was then washed 2x with PBS before incubation in 100 μl of blocking buffer (5% BSA, 0.1% Triton-X) for 1 hour at RT. Cells were incubated with primary antibody in blocking buffer overnight at 4°C. The next day wells were washed 3x with blocking buffer, before incubating with secondary antibodies for 1 hour at RT in the dark. Secondary antibodies used were Alexa-488 anti-Rabbit and Alexa 568 anti-Mouse (Invitrogen). Wells were washed 2x in blocking buffer and 1x in PBS, before incubation in 4′,6-diamidino-2-phenylindole (DAPI). 2x PBS washes were then performed and each well was kept in 100 μl PBS until imaging. Imaging was performed using an Opera Phenix Plus with a 40x objective.

### Image Analysis

Nuclei segmentation, cytoplasm detection, and fluorescence intensity measurements were performed using Harmony 4.9 software. Intensity measurements for each cell were exported from the Harmony software for analysis and visualization using RStudio 4.3. For complementation cell lines, the expression of the mutant protein was defined based on a higher HA intensity in the cytoplasm compared to the parental cell line which does not express HA-tagged proteins. Only HA-positive cells were used in the analysis for the complementation cell lines. Images for visualization were exported to ImageJ, where brightness and contrast were edited for visibility without altering pixel values. All images were processed using the same steps. Figures with visualizations were prepared using QuickFigures^66^.

### Statistics

Statistical tests on microscopy data were performed between the medians of multiple biological replicates. For comparisons between multiple conditions, paired ANOVA was used paired by the replicate. When the ANOVA showed a difference between conditions, paired t-test was used to find the difference between specific conditions. Significance threshold was defined as p<0.05.

## Data availability

The cryo-EM map was deposited to EMDB (ID: EMD-60697, https://www.ebi.ac.uk/pdbe/entry/emdb/EMD-60697) and the coordinates of the atomic structure to PDB (ID: 9IMZ, https://www.rcsb.org/structure/9IMZ). Other original data were provided in a resource data file.

## Acknowledgement

We thank the members of Song lab for helpful discussion. We thank KAIST Analysis Center for Advanced Research (KARA), Institute of Membrane Proteins (IMP), Research Resource Center at Institute of Basic Science (IBS), and Global Science Data Center (GSDC) and KREONET at Korea Institute of Science and Technology Information (KISTI) for supporting cryo-EM facilities and computing resources. The image processing package was supported from the SBgrid (www.sbgrid.org). We thank the Protein Imaging Platform at the Novo Nordisk Foundation Center for Protein Research (CPR) for assistance and support in the microscopy and image analysis facilities. This work is partially supported by grants (RS-2024-00333346, RS-2023-00266300, RS-2024-00440614 to J.S.) by National Research Foundation of Korea and Grand Challenge 30 by KAIST. T.K. is supported by Brain Korea 21 program. Research in A.G.’s laboratory was supported by the Novo Nordisk Foundation (NNF21OC0067425) and the European Research Council (CoG 724436). Research at CPR is supported by the Novo Nordisk Foundation (NNF14CC0001).

## Author contributions

J.S. and A.G. conceptualized the idea and supervised all experiments. T.J., R.C.M.F., J.Y. performed experiments. All authors reviewed the data and wrote the paper.

