## Supplementary data for "CODANIN-1 sequesters ASF1 by using a histone H3 mimic helix to regulate the histone supply"

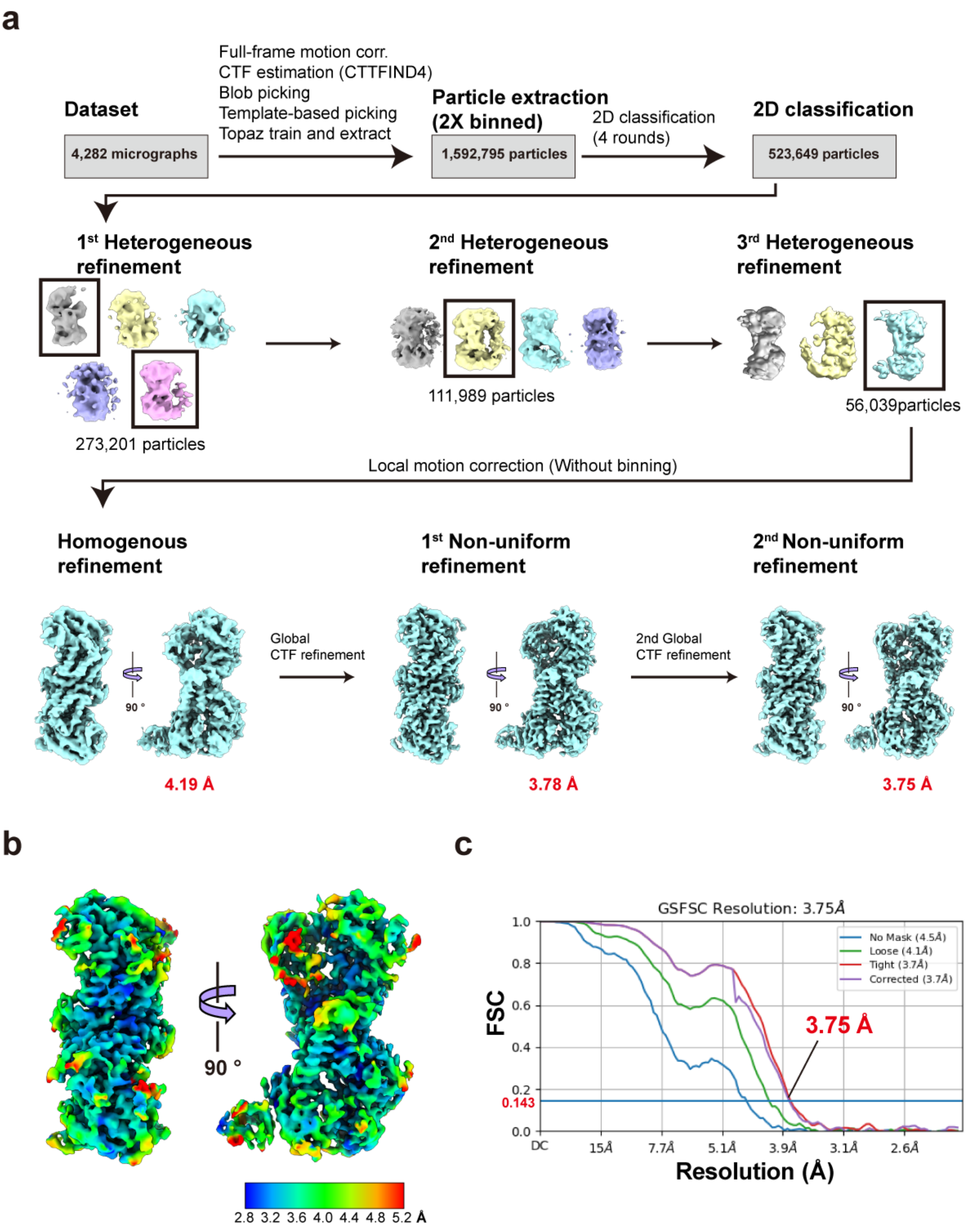

**Supplementary Fig. 1 Cryo-EM data processing.**

**a,** A flow chart of image processing to determine the cryo-EM structure of CODANIN-1 complex using CryoSPARC v4.2.1. **b,** A Local resolution map of the CODANIN-1 complex. **c,** Gold Standard Fourier Shell Curve (FSC) showing the resolution of 3.75 Å at FSC=0.143.

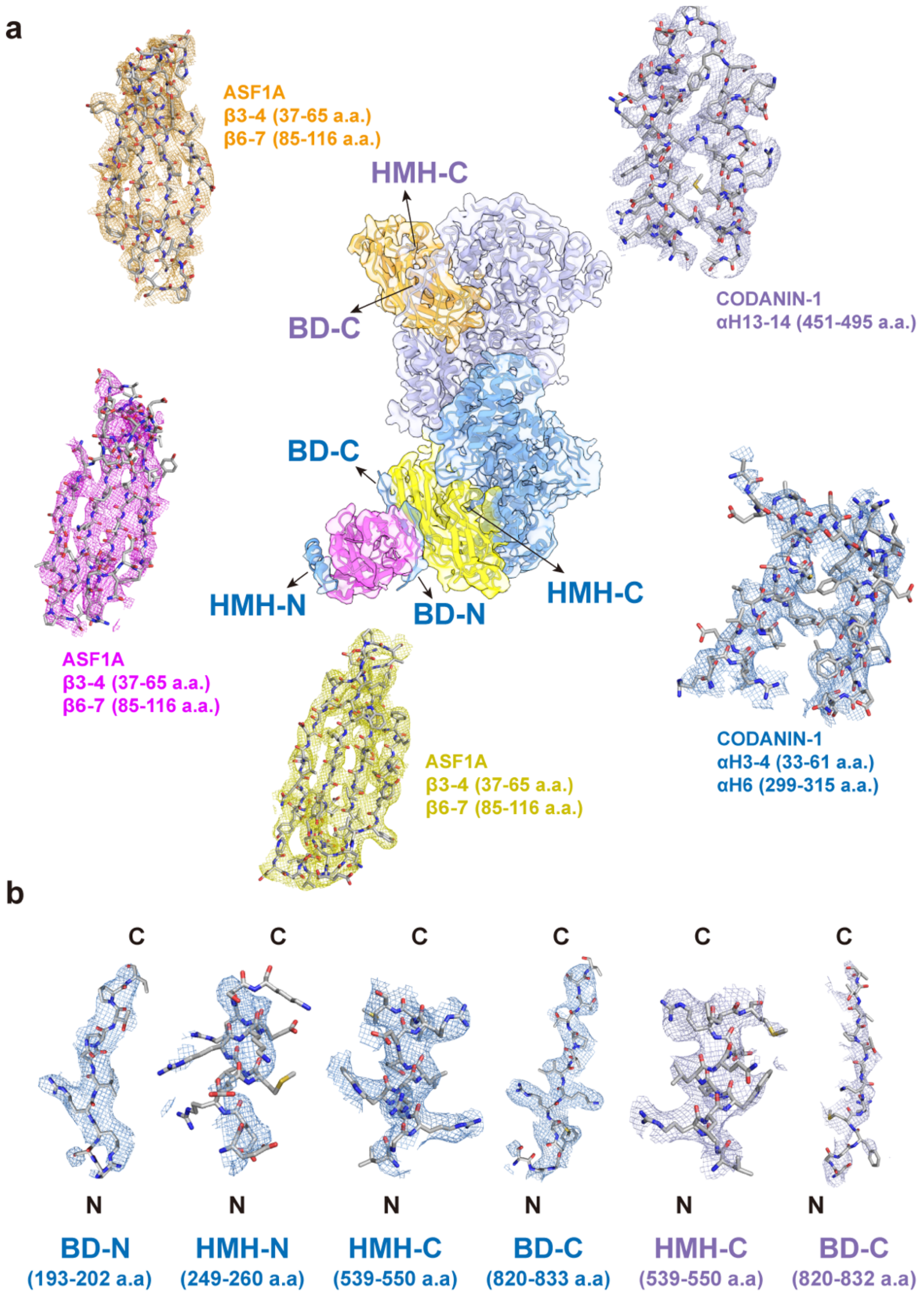

**Supplementary Fig. 2 The Cryo-EM map with a fitted atomic model.**

**a,** Representative areas of the cryo-EM map fitted with the atomic model showing the high quality of the map. Density map was drawn by isomesh command in Pymol. Map contoured at 2.0 sigma within 2.0 Å, except for the $_{dist}$ASF1A, which was contoured at 4 sigma within 2.0 Å. **b**, cryo-EM maps and fitted atomic models for each BD-N, HMH-N, HMH-C, and BD-C on CODANIN-1 chain A, as well as HMH-C and BD-C on CODANIN-1 chain B.

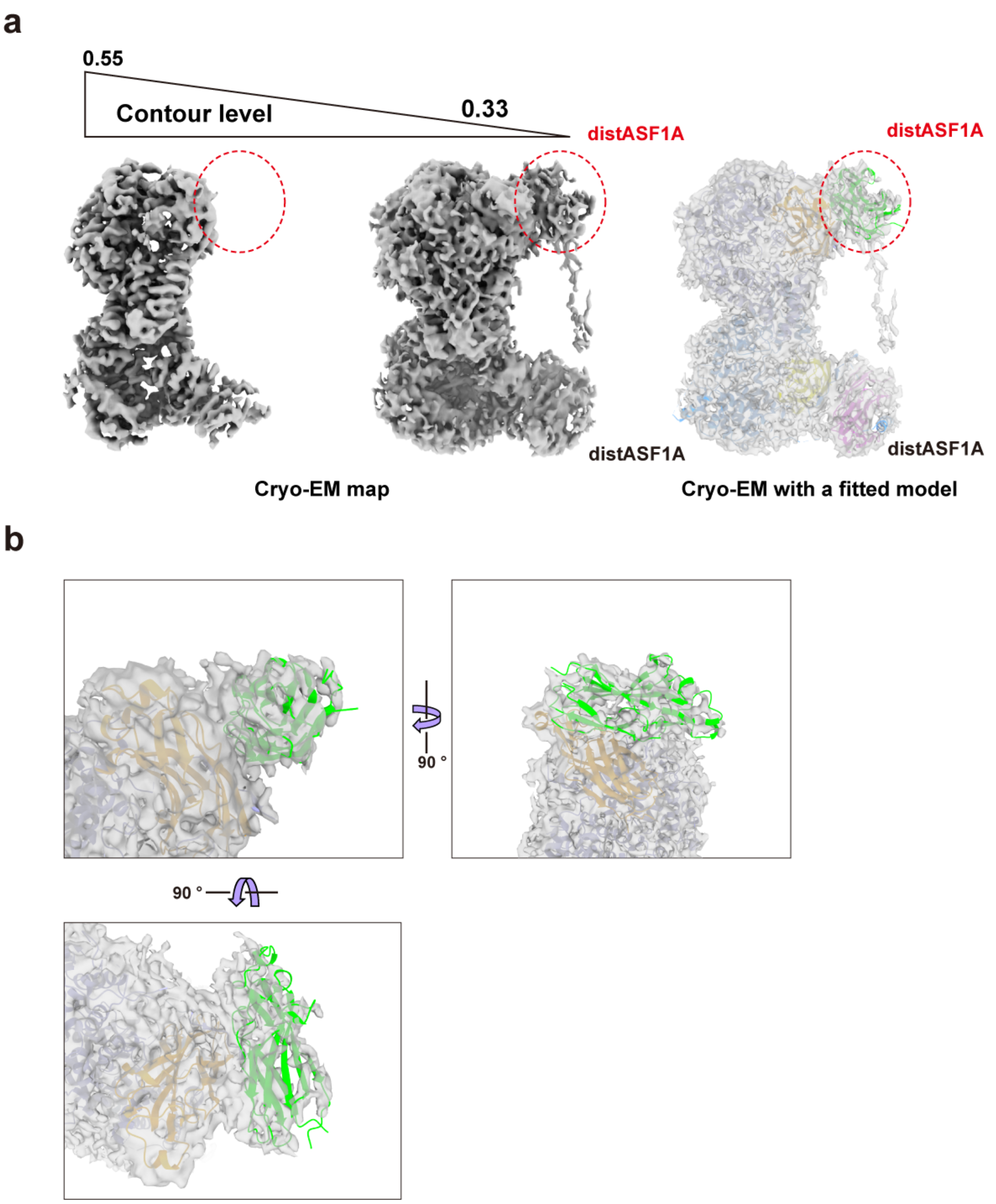

**Supplementary Fig. 3 Cryo-EM map of CODANIN-1 ASF1A complex in 2:4 stoichiometry.** **a**, The cryo-EM map is displayed at contour levels of 0.55 and 0.33. The low-contoured-level density map was fitted with atomic models for the CODANIN-1 complex and distal ASF1A molecules. An additional volume, representing a distal ASF1A, is highlighted with a red dashed circle. **b**, An enlarged view of the additional volume fitted with the distal ASF1A model (colored green).

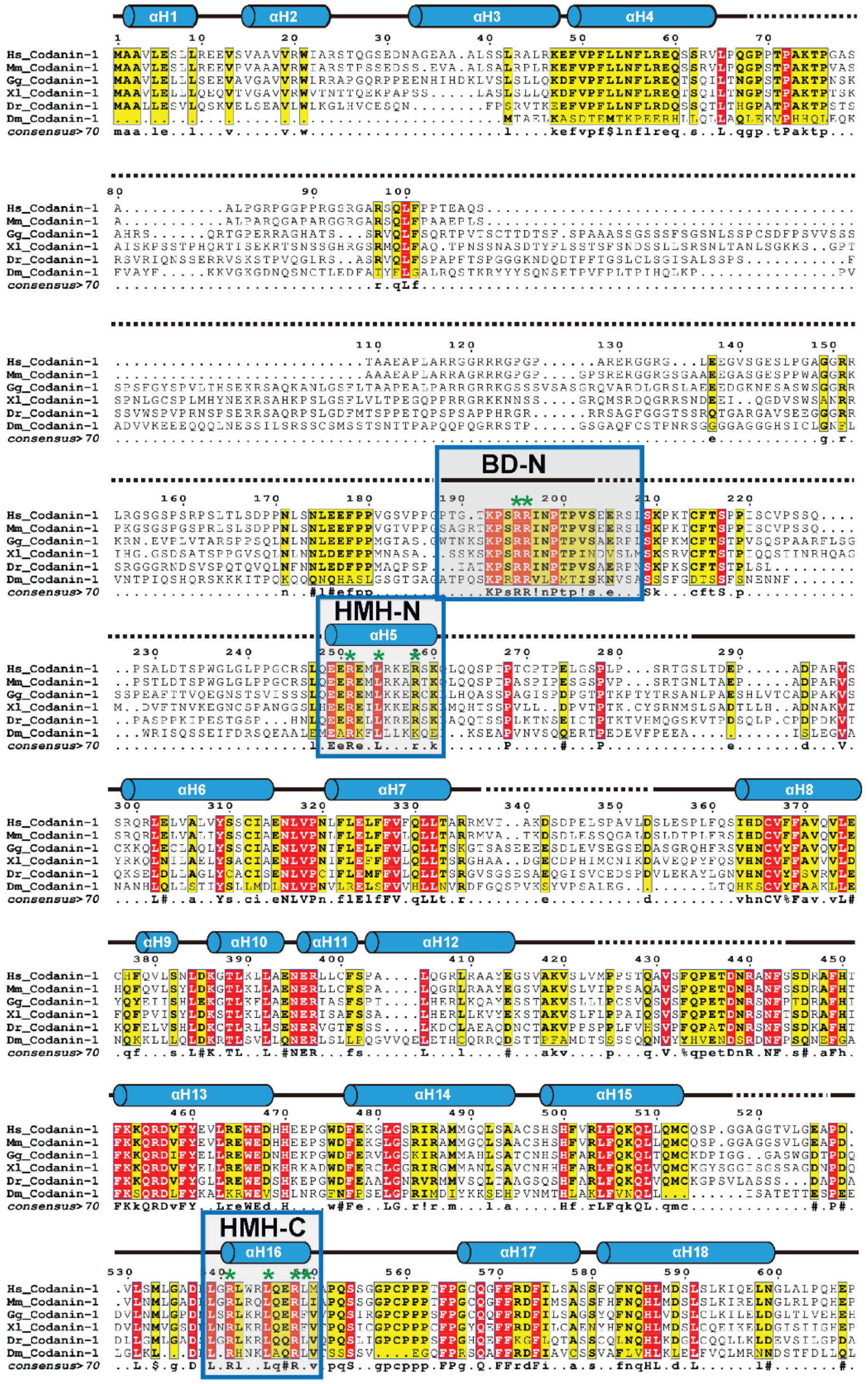
αH1
αH2
αH3
αH4
BD-N
HMH-N
αH5
αH6
αH7
αH8
αH9
αH10
αH11
αH12
αH13
αH14
αH15
HMH-C
αH16
αH17
αH18
Hs_Codanin-1
Mm_Codanin-1
Gg_Codanin-1
Xl_Codanin-1
Dr_Codanin-1
Dm_Codanin-1
consensus>70

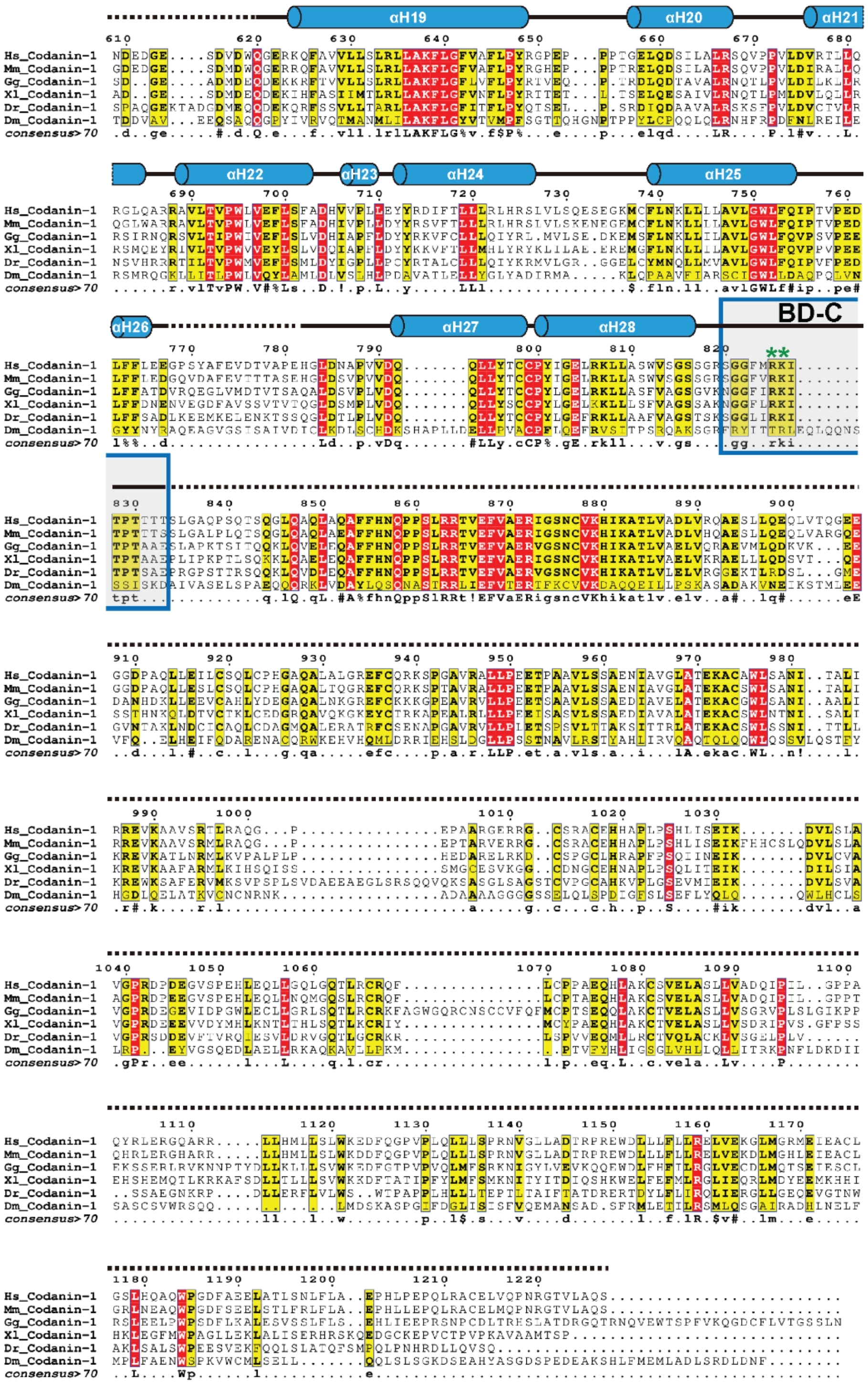

**Supplementary Fig. 4 Sequence alignment of CODANIN-1.**

A sequence alignment of CODANIN-1 among human (Hs), mouse (Mm), chicken (Gg), xenopus (Xl), zebrafish (Dr) and fly (Dm). The secondary structural elements from the cryo-EM structure are shown above the sequence. Disordered areas are indicated with dotted lines. The B-domains and HMHs are highly conserved across various species (highlighted by boxes). The residues involved in ASF1 interaction at HMH and B-domain, are marked with asterisks.

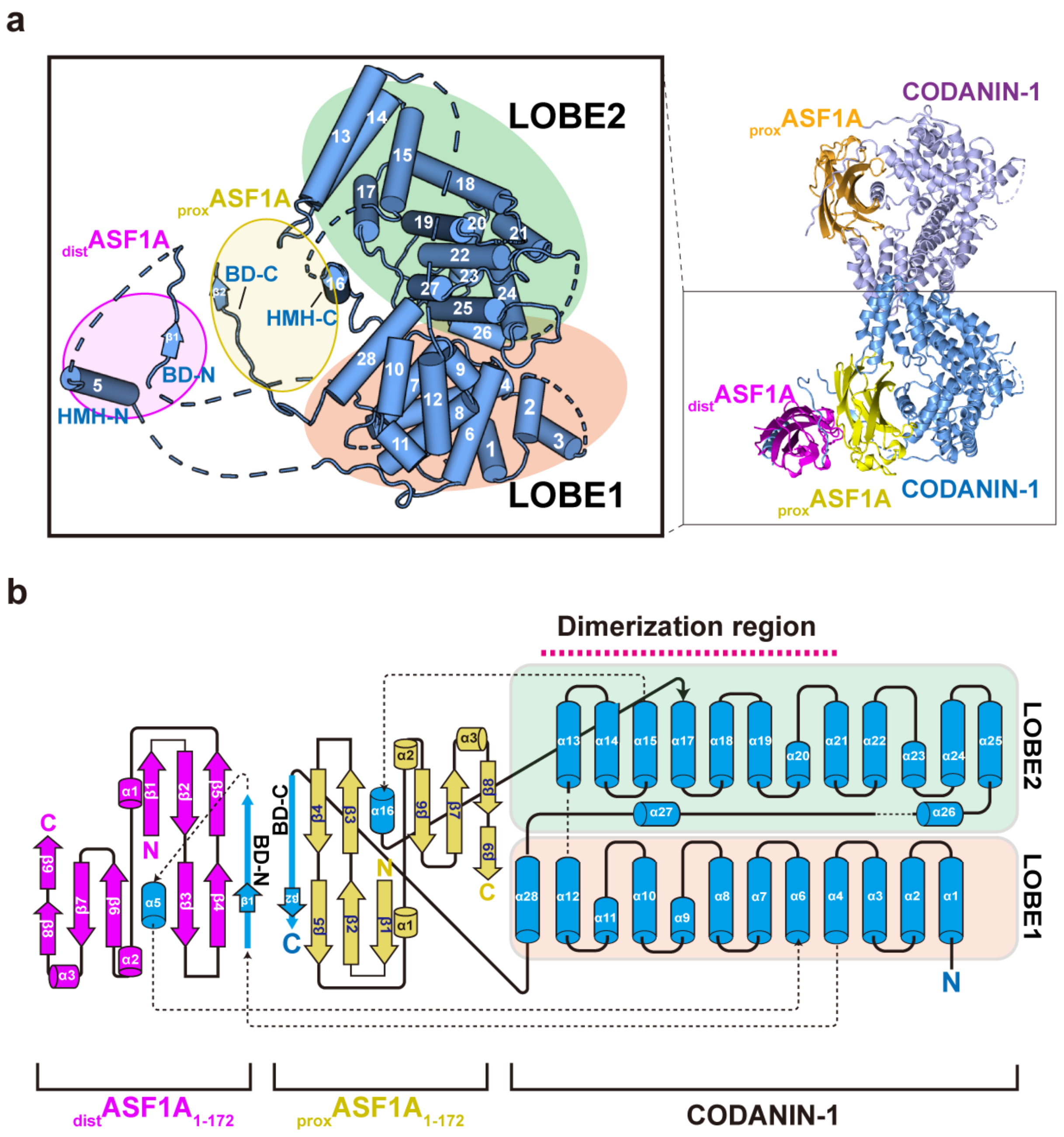

**Supplementary Fig. 5 The domain organization of CODANIN-1 monomer and ASF1A complex.** **a,** CODANIN-1 is composed of Lobe1 and Lobe2. Each of 28 α-helices and two β-strands identified in the cryo-EM structure is labeled on the cartoon representation of the structure. The position of ASF1A was marked as an ellipse in the box. **b,** A topology diagram of cryo-EM structure of CODANIN-1 and ASF1A. Cylinders represent α-helix, arrows for β-strands thick lines for loops, and the disordered loops are drawn in dotted lines. The B-domain of CODANIN-1 was highlighted with a thick blue arrow for emphasis. The dimerization interface of CODANIN-1 is marked red dotted line.

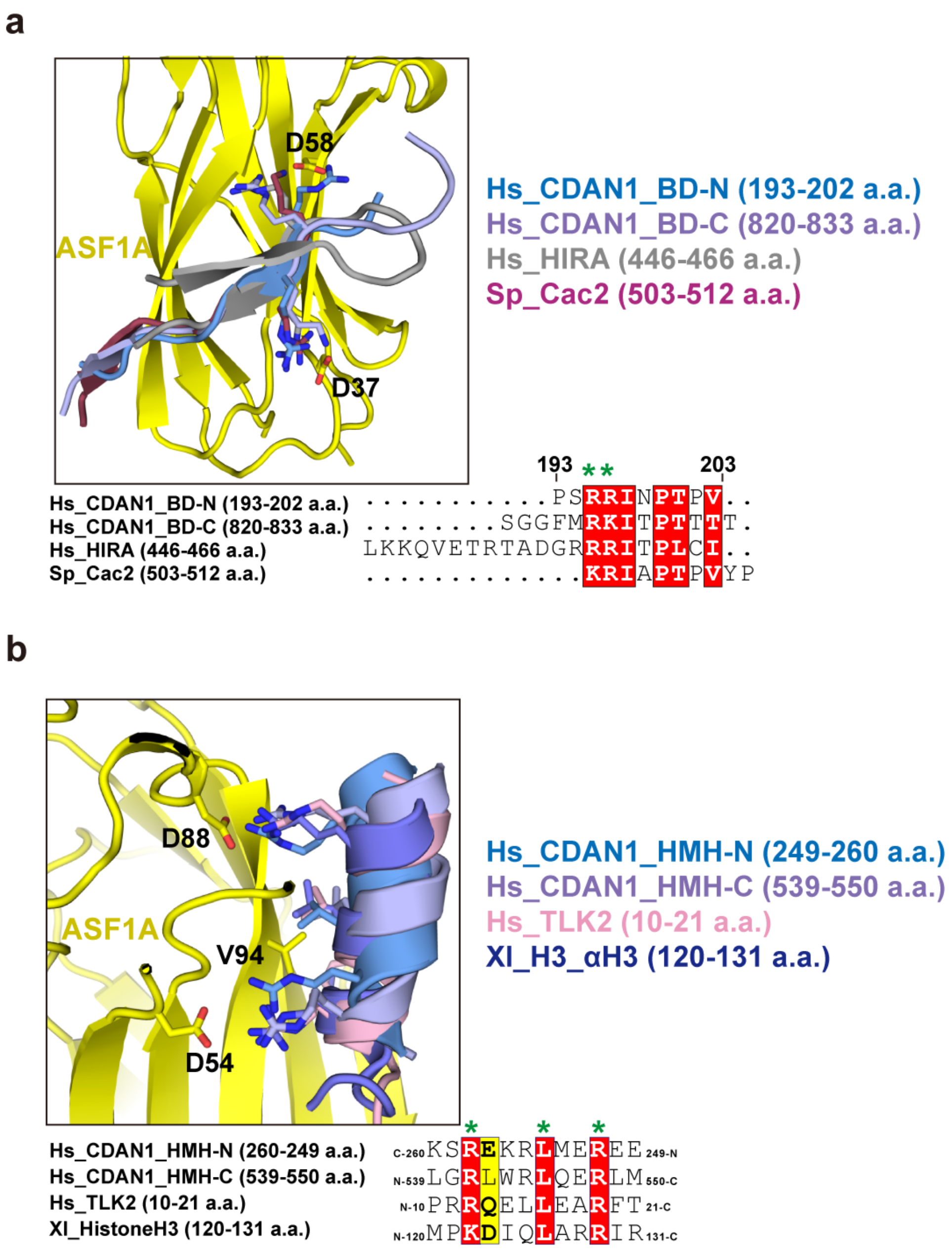

**Supplementary Fig. 6 Conservation of the B-domain and HMH.**

**a**, Superimposing the B-domains of CODANIN-1 (BD-N in pale blue and BD-C in pale purple), HIRA (PDB 2I32, shown in grey), and Schizosaccharomyces pombe (Sp) CAF-1 subunit Cac2 (PDB 2Z3F, shown in violet). A sequence alignment among the B-domains from CODANIN-1, HIRA and CAF-1 (below). The residues involving ASF1 interaction are marked with asterisks. **b,** Superimposing the HMHs of CODANIN-1 (HMH-N in pale blue and HMH-C in pale purple), a histone mimic helix of TKL2 (PDB 7LO0 in pink) and αH3 helix of histone H3 (PDB 2IO5, in deepblue). A sequence alignment of HMH helix of CODANIN-1 and TLK2 and histone H3 αH3 (below). The residues involving ASF1 interaction are marked with asterisks.

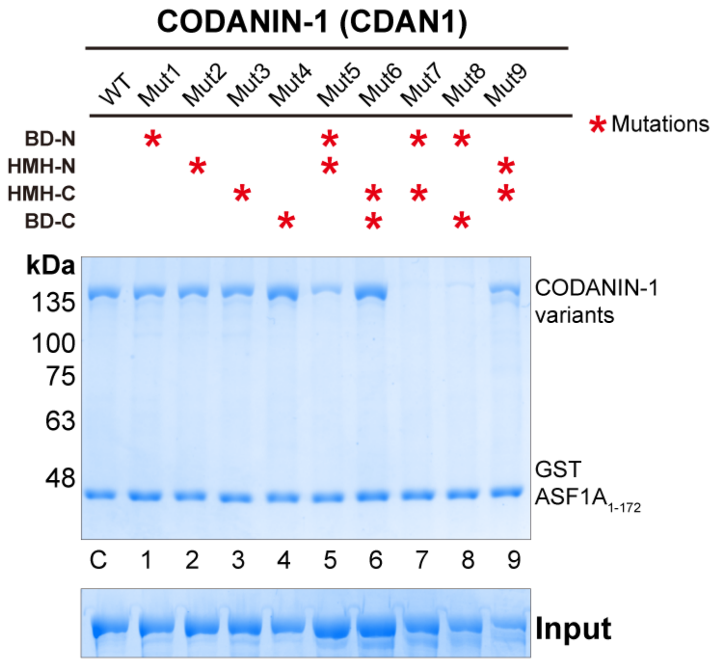

**Supplementary Fig. 7 GST pulldown experiments for the interaction between CODANIN-1 and ASF1A.**

An SDS-PAGE gel of the GST pulldown experiments showing the binding of full-length CODANIN-1 or its mutants to GST-ASF1A. A table summarizing the mutation motifs for each mutant is provided above the gel. The mutants correspond to those shown in Fig. 3c. The input gel in the lower panel.

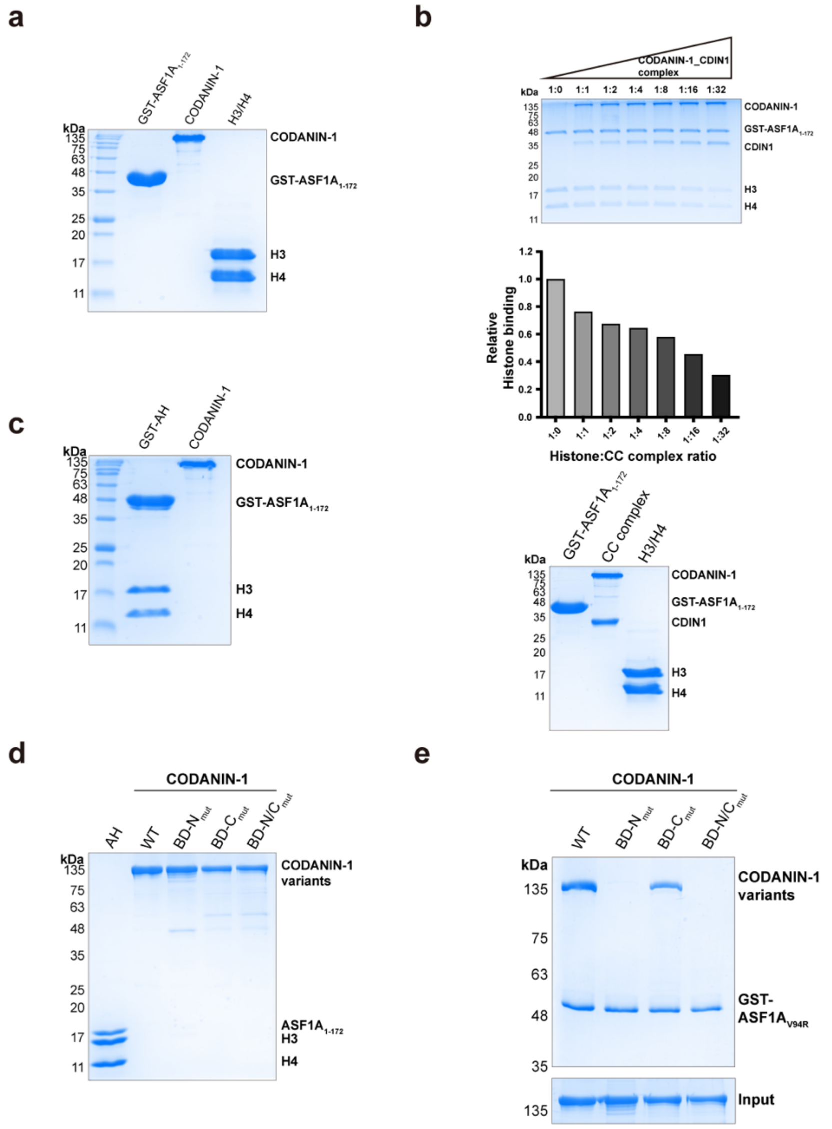

**Supplementary Fig. 8 Interaction between CODANIN-1 and ASF1A histone complex**

**a,** An SDS-PAGE gel of the inputs for the competition assay for Fig. 4a. **b,** An SDS-PAGE gel demonstrating the competitive relationship between the CODANIN-1_CDIN1 complex and histone H3-H4 (top panel). The ratio of histone H3-H4 to CODANIN-1_CDIN1 complex is indicated on each gel lane. Quantification of the relative intensity of histone H3-H4 band to GST-ASF1A band intensity (n=1 independent experiment, middle panel). The values were normalized against histone H3-H4 bands in the absence of CODANIN-1 (1:0 ratio). An SDS-PAGE gel of the inputs for the competition assay between the CODANIN-1_CDIN1 complex and histone H3-H4 (lower panel). **c,** An SDS-PAGE gel of the inputs for the competition assay for Fig. 4b. **d,** An SDS-PAGE gel of the inputs for the FLAG-pulldown experiments for Fig. 4c. **e,** An SDS-PAGE gel from the GST-pulldown experiment showing the binding of full-length CODANIN-1 and its B-domain mutants (BD-$N_{mut}$, BD-$C_{mut}$, and BD-N/$C_{mut}$) to the GST-ASF1$A_{V94R}$ mutant, which has disrupted HMH binding. The input gel for CODANIN-1 samples is shown in the lower panel.

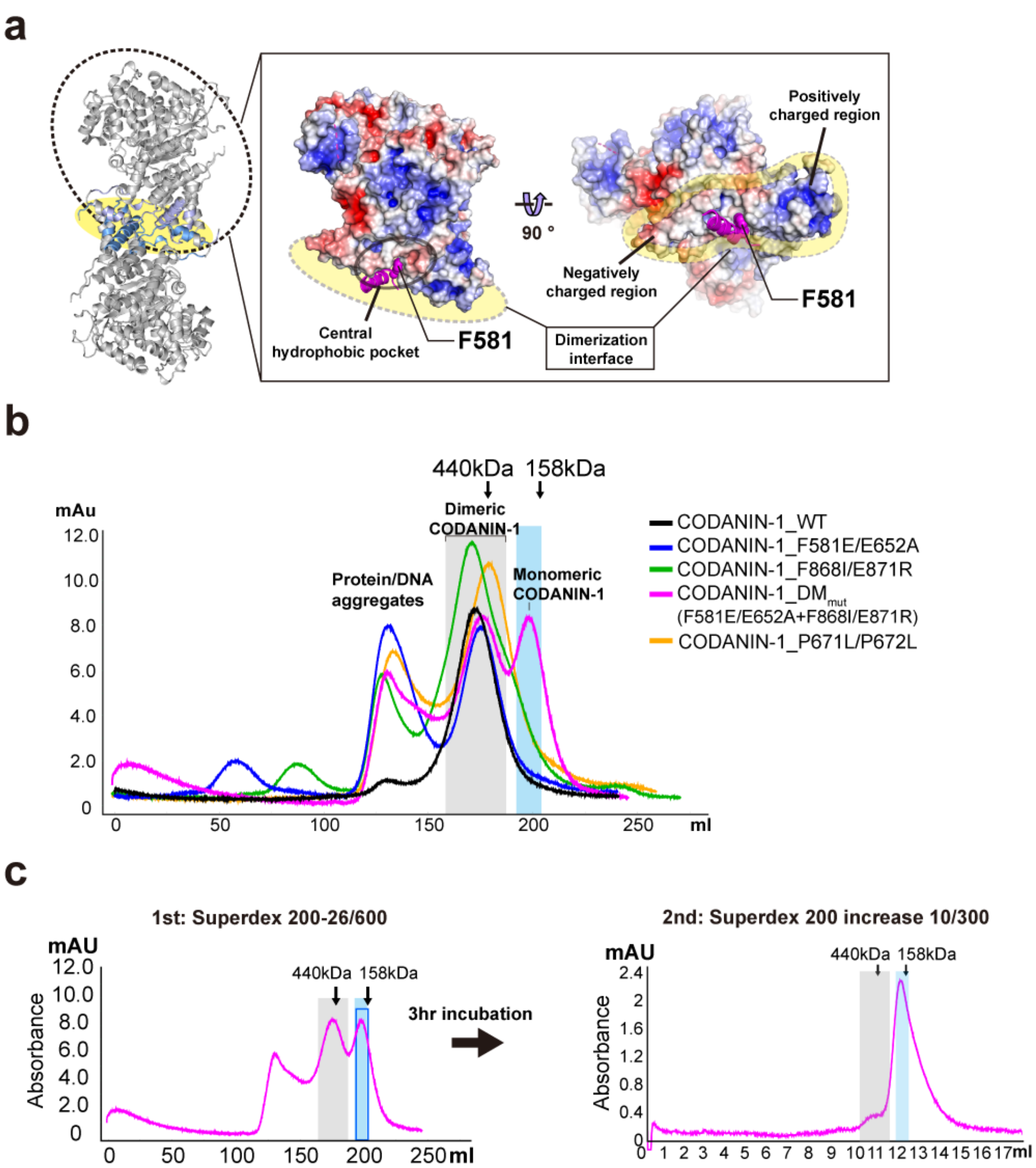

**Supplementary Fig. 9 Dimerization mutant of CODANIN-1**

**a,** Electrostatic surface representation of the dimer interface of CODANIN-1. F581 is inserted into the hydrophobic pocket formed from the other CODANIN-1. **b,** Size-exclusion chromatograms of wild type CODANIN-1 (Black) and the mutants: F581E/E652A (Blue), F868I/E871R (Green), $DM_{mut}$ (F581E/E652A+F868I/E871R, Magenta), and P671L/P672L (Orange). **c,** Series of size-exclusion chromatograms for $DM_{mut}$. The first chromatography was conducted using a HiLoad 26/600 Superdex 200 pg column (left), and the second chromatography was performed using a Superdex200 increase 10/300 GL column (right). The grey box represents the retention volume of the dimeric CODANIN-1, while the blue box indicates monomeric CODANIN-1.

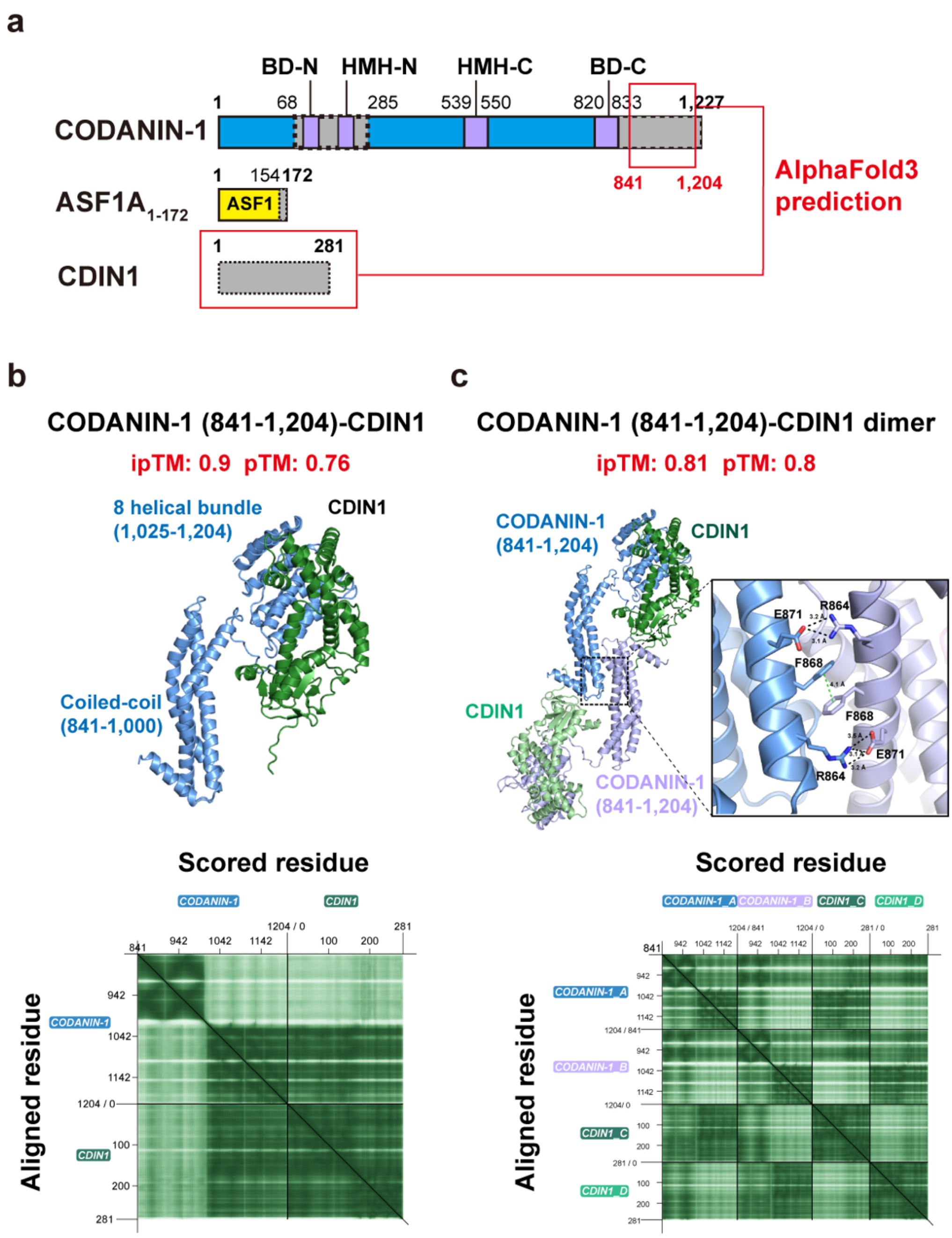

**Supplementary Fig. 10 AlphaFold2 predicted model of the C-terminal region of CODANIN-1 and CDIN1 complex.**

**a,** a schematic view of CODANIN-1 complex. Boxed regions represent AlphaFold2 predicted regions: CODANIN-1 C-terminal (841-1,204) and full-length CDIN1. **b,** AlphaFold3 predicted model and the Predicted Aligned Error (PAE) plot of CODANIN-1 (841-1,204) and CDIN1. **c,** AlphaFold3 predicted dimer model and the PAE plot for CODANIN-1 (841-1,204) and CDIN1. The dimeric interaction between the C-terminal regions is illustrated in the box. PAE plot was drawn by PAE Viewer[1].

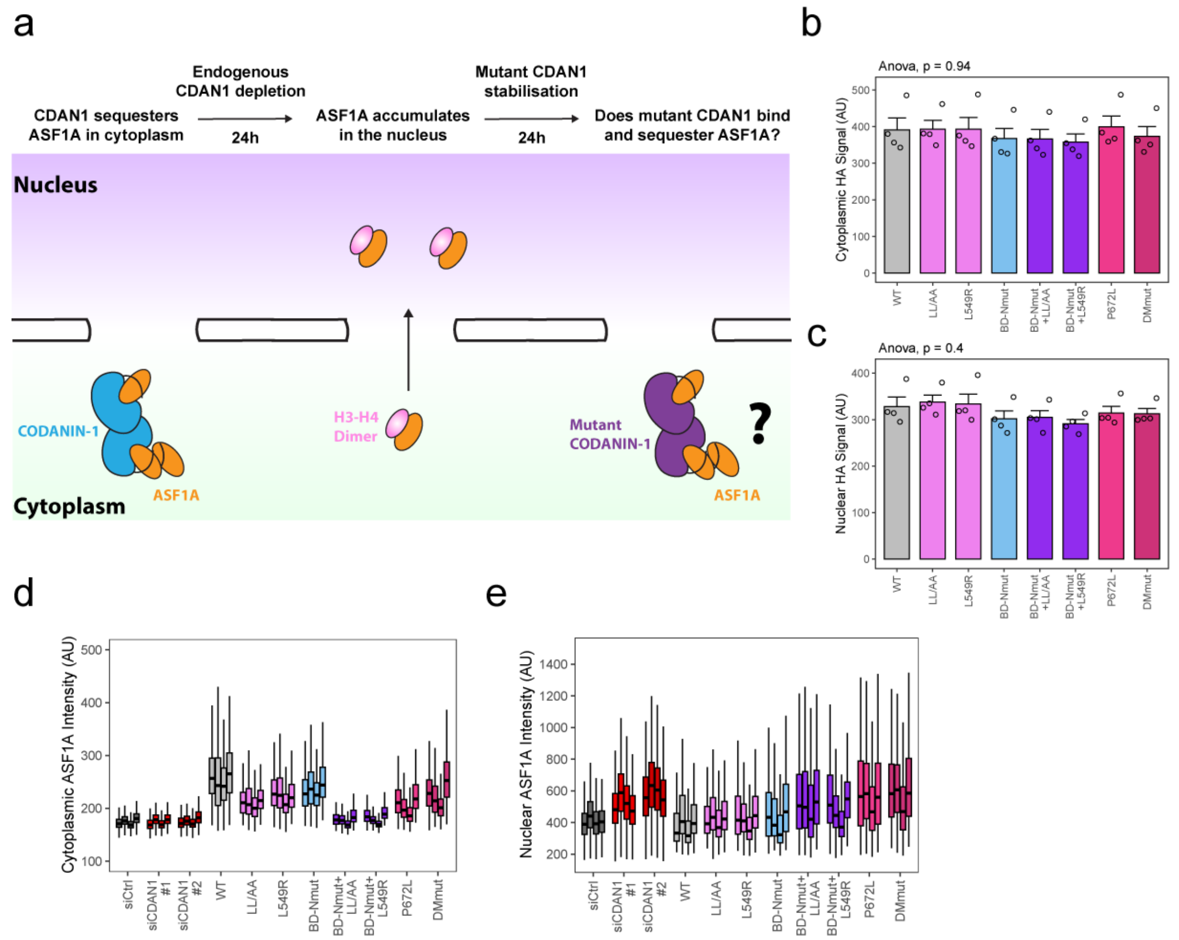

**Supplementary Fig. 11 In-cell analysis of the dual-binding mode of CODANIN-1**

**a,** Schematic of the complementation system. Removal of CDAN1 by 24 hours of siRNA knockdown leads to ASF1A translocation to the nucleus. Reintroduction of CDAN1 mutant CDAN1 construct by 24 hours of expression induction by doxycycline, and Shield1 mediated inhibition of its degradation allows assessment of mutant function. To test for ASF1A binding ability we image ASF1A localization to the nucleus, as CODANIN-1 that can bind ASF1A will cause ASF1A localization to the cytoplasm. **b,** Median cytoplasmic HA intensity of the 4 microscopy biological replicates for each induced cell line. Only cells expressing HA above background levels were considered. Each point is the median of a single biological replicate for cells induced for the indicated CODANIN-1 construct, and ANOVA paired by replicate was used to test for differences between conditions. **c,** Median nuclear HA intensity of the 4 microscopy biological replicates for each induced cell line. Only cells expressing HA above background levels were considered. Each point is the median of a single biological replicate for cells induced for the indicated CODANIN-1 construct, and ANOVA paired by replicate was used to test for differences between conditions. **d,** Boxplot of the distribution of cytoplasmic ASF1A intensity from each biological replicate in microscopy experiments shown in Figure 6c-g. Each box represents a distinct biological replicate. **e,** Boxplot of ASF1A nuclear intensity in each biological replicate shown in Figure 6c-g.

**Supplementary Table 1.** Antibodies used in this study. WB: western blot, IF: immunofluorescence.

| Target | Source | Catalogue No. | Application | Dilution |
|---|---|---|---|---|
| CDAN1 | abcam | ab204392 (Lot GR11473) | WB | 1:1000 |
| α-TUBULIN | abcam | ab6160 (lot 1011061-1) | WB | 1:5000 |
| HA | BioLegend | 901513 (clone 16B12) | WB+IF | 1:1000 |
| ASF1A | Cell Signaling | 2990S (clone C6E10) | IF | 1:100 |
| Anti-Rabbit Alexa 488 | Invitrogen | A-11034 | IF | 1:1000 |
| Anti-Mouse Alexa 568 | Invitrogen | A-11031 | IF | 1:1000 |
| IRDye 800CW anti-Rabbit IgG | LiCor-Bio | 926-32211 | WB | 1:10000 |
| IRDye 800CW anti-Rat IgG | LiCor-Bio | 926-32219 | WB | 1:10000 |
| IRDye 700CW anti-Mouse IgG | LiCor-Bio | 926-68070 | WB | 1:10000 |

**Supplementary Table 2.** Plasmids used and generated in this study.

| Plasmid | Source | Reference |
|---|---|---|
| pcDNA5/FRT/TO-FLAG-HA-CDAN1 | Ask et al. 2012 (Ask *et al.*, 2012) | |
| pcDNA5/FRT/TO-FLAG-HA-DD-CDAN1 | This study | |
| pcDNA5/FRT/TO-FLAG-HA-DD-CDAN1-L545A_L549A | This study | |
| pcDNA5/FRT/TO-FLAG-HA-DD-CDAN1-L549R | This study | |
| pcDNA5/FRT/TO-FLAG-HA-DD-CDAN1-R195A_R196A | This study | |
| pcDNA5/FRT/TO-FLAG-HA-DD-CDAN1-R195A_R196A_L545A_L549A | This study | |
| pcDNA5/FRT/TO-FLAG-HA-DD-CDAN1-R195A_R196A_L549R | This study | |
| pcDNA5/FRT/TO-FLAG-HA-DD-CDAN1-P672L | This study | |
| pcDNA5/FRT/TO-FLAG-HA-DD-CDAN1-F581E_E652A_F868I_E871R | This study | |
| pOG44 | ThermoFisher | V600520 |
| pFastBac1/CDAN1-6xHis | This study | |
| pFastBac1/CDIN1 | This study | |
| pET21a/ASF1A (1-172) | This study | |
| pFastBac1/CDAN1-FLAG | This study | |
| pFastBac1/CDAN1_BD-$N_{mut}$-FLAG | This study | |
| pFastBac1/CDAN1_HMH-$N_{mut}$-FLAG | This study | |
| pFastBac1/CDAN1_HMH-$C_{mut}$-FLAG | This study | |
| pFastBac1/CDAN1_BD-$C_{mut}$-FLAG | This study | |
| pFastBac1/CDAN1_BD-$N_{mut}$ / HMH-$N_{mut}$-FLAG | This study | |
| pFastBac1/CDAN1_HMH-$C_{mut}$ / BD-$C_{mut}$-FLAG | This study | |
| pFastBac1/CDAN1_BD-$N_{mut}$ / HMH-$C_{mut}$-FLAG | This study | |
| pFastBac1/CDAN1_BD-$N_{mut}$ / BD-$C_{mut}$-FLAG | This study | |
| pFastBac1/CDAN1_HMH-$N_{mut}$ / HMH-$C_{mut}$-FLAG | This study | |
| pGEX-4T-1/ASF1A (1-172) | This study | |
| pGEX-4T-1/ASF1A (1-172)_D88A | This study | |

| | |
|---|---|
| pGEX-4T-1/ASF1A (1-172)_D54A | This study |
| pGEX-4T-1/ASF1A (1-172)_V94R | This study |
| pGEX-4T-1/ASF1A (1-172)_E36A/D37A | This study |
| pGEX-4T-1/ASF1A (1-172)_D88A_E36A/D37A | This study |
| pGEX-4T-1/ASF1A (1-172)_D54A_E36A/D37A | This study |
| pGEX-4T-1/ASF1A (1-172)_V94R_E36A/D37A | This study |
| pGEX-4T-1/ASF1A (193-202) | This study |
| pGEX-4T-1/ASF1A (249-260) | This study |
| pGEX-4T-1/ASF1A (539-550) | This study |
| pGEX-4T-1/ASF1A (820-833) | This study |

**Supplementary Table 3**. Primers used in this study for cloning and genotyping.

| Primer | Sequence |
|---|---|
| CDAN1 genotyping 1 | ATGGCTGCTGTGCTGGAAAG |
| CDAN1 genotyping 2 | TGCAGGAAGAAAGAGAGATGC |
| CDAN1 genotyping 3 | CGCCTGTAGCCACTCTCATTTC |
| CDAN1 genotyping 4 | ATGTGCTTCCTGAACAAACTGC |
| CDAN1 genotyping 5 | TGATCAGACGGGAAGTGAAAGC |
| FLAG-HA-DD genotyping forward | CAACTCCGCCCCATTGACGCAA |
| FLAG-HA-DD genotyping reverse | AGGTCCCTGAGGCAGCACTCTG |
| B domain forward | CGCAGCAATCAACCCCACCCCCGTG |
| B domain reverse | CTGGGCTTTGTGCCGGTAGGTCC |
| HMH1 forward | CCTGCGCCCGCCACAGTCT |
| HMH1 reverse | AACGGGCGATGGCTCCTCAGTCTAG |
| HMH2 forward | GCAGGAACGGCGCATGGCTCCTC |
| HMH2 reverse | AGCCGCCACAGTCTG |
| F581E forward | CGCCAGCAGCGAACAGTTCAACC |
| F581E reverse | CTCAGGATGAAATCCCG |
| E652A forward | CAGAGGACCCGCGCCTCCTCCAA |
| E652A reverse | TATGGCAGAAAGGCCACGAAG |
| P672L forward | CCAGGTGCCACTGGTGCTGGATG |
| P672L reverse | CTTCTCAGGGCCAGG |
| F861I_E871R forward | CACAATTTCCACGGTTCTCCGCAG |
| F861I_E871R reverse | GCCAGGAGAATCGGCAGCAAC |
| DD megaprimer forward | CGACTACGCGGGATCCGGAGTGCAGGTGGAAACCATC |
| DD megaprimer reverse | CCAGCACAGCAGCCATGATATCTTCCGGTTTTAGAAGCTCCAC |

| | |
|---|---|
| CDAN1 BD-Nmut forward | CCGACCGGCACCAAACCGAGTGCTGCTATTAATCCGACACCGG |
| CDAN1 BD-Nmut reverse | CCGGTGTCGGATTAATAGCAGCACTCGGTTTGGTGCCGGTCGG |
| CDAN1 HMH-Nmut forward | CTGCAAGAGGAACGTGAAATGCGGCGCAAAGAAC |
| CDAN1 HMH-Nmut reverse | GTTCTTTGCGCCGCATTTCACGTTCCTCTTGCAG |
| CDAN1 HMH-Cmut forward | GTCGCCTGTGGCGTGCGCAAGAACGTGCGATGGCACCGCAGAGC |
| CDAN1 HMH-Cmut reverse | GCTCTGCGGTGCCATCGCACGTTCTTGCGCACGCCACAGGCGAC |
| CDAN1 BD-Cmut forward | GCGGTCGTAGCGGTGGTTTTATGGCTGCAATTACCCCGACCACAACC |
| CDAN1 BD-Cmut reverse | GGTTGTGGTCGGGGTAATTGCAGCCATAAAACCACCGCTACGACCGC |
| CDAN1 F581E forward | CTGAGCGCAAGCAGCGAACAGTTTAATCAGCATCTGATGG |
| CDAN1 F581E reverse | CCATCAGATGCTGATTAAACTGTTCGCTGCTTGCGCTCAG |
| CDAN1 E652A forward | GTATCGTGGTCCGGCACCGCCACCGACCGGTGAAGCGCAGGATAGCATTC |
| CDAN1 E652A reverse | GAATGCTATCCTGCGCTTCACCGGTCGGTGGCGGTGCCGGACCACGATAC |
| CDAN1 F868I forward | CGAGTCTGCGTCGTACCGTTGAAATTGTTGC |
| CDAN1 F868I reverse | GCAACAATTTCAACGGTACGACGCAGACTCG |
| CDAN1 E871R forward | GCGTCGTACCGTTGAATTTGTTGCACGACGTATTGGTAGC |
| CDAN1 E871R reverse | GCTACCAATACGTCGTGCAACAAATTCAACGGTACGACGC |

| | |
|---|---|
| ASF1A V94R forward | GCAGATGCAGTAGGCGTAACTCGTGTGCTAATTACTTGTACC |
| ASF1A V94R reverse | GGTACAAGTAATTAGCACACGAGTTACGCCTACTGCATCTGC |
| ASF1A E36A/D37A forward | GCATCGAGGACCTGTCTGCAGCCTTGGAATGGAAAATTATC |
| ASF1A E36A/D37A reverse | GATAATTTTCCATTCCAAGGCTGCAGACAGGTCCTCGATGC |
| ASF1A D88A forward | GGACTCATTCCAGATGCAGCTGCAGTAGGCG |
| ASF1A D88A reverse | CGCCTACTGCAGCTGCATCTGGAATGAGTCC |
| ASF1A D54A forward | GGGCTCTGCAGAAAGTGAAGAATACGCTCAAGTTTTAGACTCTG |
| ASF1A D54A reverse | CAGAGTCTAAAACTTGAGCGTATTCTTCACTTTCTGCAGAGCCC |

**Supplementary Table 4**. Cell lines used and generated in this study.

| Cell Lines | Description | Source |
|---|---|---|
| Spodoptera frugiperda 9 | Cells used for CDAN1 expression in cryo-EM study and in vitro biochemical assays. | Thermo Fisher Scientific |
| U2OS_Flp In/TREX | Cell line for integration of tetracycline-inducible constructs by FlpIn. | Gift from J. Nilsson lab |
| U2OS_FlpIn_DD-FLAG-HA-CDAN1_WT | Tetracycline-inducible expression of wild type CDAN1 with Shield1-inducible stabilisation | This study |
| U2OS_FlpIn_DD-FLAG-HA-CDAN1_L549R | Tetracycline-inducible expression of L549R CDAN1 with Shield1-inducible stabilisation | This study |
| U2OS_FlpIn_DD-FLAG-HA-CDAN1_ L545A_L549A | Tetracycline-inducible expression of LL/AA CDAN1 with Shield1-inducible stabilisation | This study |
| U2OS_FlpIn_DD-FLAG-HA-CDAN1_WT_R195A_R196A | Tetracycline-inducible expression of $B_{mut}$ CDAN1 with Shield1-inducible stabilisation | This study |

| | | |
|---|---|---|
| U2OS_FlpIn_DD-FLAG-HA-CDAN1_WT_R195A_R196A_L545A_L549A | Tetracycline-inducible expression of $B_{mut}$+LL/AA CDAN1 with Shield1-inducible stabilisation | This study |
| U2OS_FlpIn_DD-FLAG-HA-CDAN1_R195A_R196A_L549R | Tetracycline-inducible expression of $B_{mut}$+L549R CDAN1 with Shield1-inducible stabilisation | This study |
| U2OS_FlpIn_DD-FLAG-HA-CDAN1_P672L | Tetracycline-inducible expression of CDAN1 with P672L mutation with Shield1-inducible stabilisation | This study |
| U2OS_FlpIn_DD-FLAG-HA-CDAN1_F581E_E652A_F868I_E871R | Tetracycline-inducible expression of DMmut CDAN1 with Shield1-inducible stabilisation | This study |
